# Multi-substrate DNA stable isotope probing reveals guild structure in bacterially mediated soil carbon cycling

**DOI:** 10.1101/2021.03.19.436178

**Authors:** Samuel E. Barnett, Nicholas D. Youngblut, Chantal N. Koechli, Daniel H. Buckley

**Author notes:** Co-first authors (Samuel E. Barnett, Nicholas D. Youngblut). Corresponding author: Daniel H. Buckley. **Author contributions:** NDY and DHB conceived of the study, NDY and CNK conducted the experiment and collected all the data, NDY and DHB analyzed the data, NDY, SEB, and DHB wrote the manuscript. All authors were involved in editing and reviewing manuscript. **Competing interests:** The authors declare no competing interests.

## Abstract

Soil microorganisms determine the fate of soil organic matter (SOM), and their activities comprise a major component of the global carbon (C) cycle. We employed a multi-substrate DNA-stable isotope probing experiment to track bacterial assimilation of C derived from distinct sources that varied in bioavailability. This approach allowed us to measure microbial contributions to SOM processing by measuring the C assimilation dynamics of diverse microorganisms as they interact within soil. We identified and tracked 1,286 bacterial taxa that assimilated ^13^C in an agricultural soil over a period of 48 days. Overall ^13^C-assimilation dynamics of bacterial taxa, defined by the source and timing of the ^13^C they assimilated, exhibited low phylogenetic conservation. We identified bacterial guilds comprised of taxa that had similar ^13^C assimilation dynamics. We show that C source bioavailability explained significant variation in both C mineralization dynamics and guild structure. In addition, guild structure explained significant variation in bacterial growth dynamics. We demonstrate that the observed guild structure is consistent with predictions made by bacterial life history theory. We also demonstrate that the guild structure explains significant variation in the biogeographical distribution of bacteria at continental and global scales. We interpret these findings in the context of bacterial life history strategies and their relationship to terrestrial C-cycling.

## Introduction

Soil organic matter (SOM) represents the largest terrestrial pool of organic carbon (C) on Earth (1), and SOM dynamics have major impacts on the global C-cycle. SOM is sensitive to land management (2–4) and climate change (5), and SOM pools influence soil health (6, 7), agricultural productivity (8), and climate stability (6, 9). However, the mechanisms that promote SOM persistence and loss remain uncertain and this introduces variability into global C-cycle models (10–13). Microorganisms are dominant drivers of SOM decomposition, stabilization, and mineralization (14), and C-cycle models increasingly seek to incorporate microbial traits to improve predictions of SOM dynamics (15, 16). For example, microbially explicit C-cycle models, such as MEND (17), MIMICS (18), and CORPSE (19), improve prediction of C-cycle dynamics when compared to models that ignore microbial processing (20).

Soil microbes play a critical role in governing global C flux, but the microbial mechanisms that determine SOM dynamics remain poorly characterized (13). Microbial contributions to soil C-cycling are typically determined in the aggregate (*e.g*., soil respiration) without providing mechanistic insight into processes of individual microbes. Determining the characteristics of individual microbes typically requires laboratory cultivation, but most soil microbes remain uncultivated and poorly characterized (21–23). In the absence of cultivation, the characteristics of uncultivated microbes are often inferred by assuming phylogenetic conservation with cultivated representatives (24–26). However, phylogenetic conservation varies dramatically among microbial traits and functionalities (27), and many soil microbes lack closely related cultivated isolates (28). Microbial characteristics are also often inferred through metagenomic analyses that indicate the frequency of specific gene families and metabolic pathways in the environment (29, 30).

Inferences based on phylogenetic conservatism and metagenomic analyses, however, predict ‘potential metabolism’, but this information cannot currently predict ecological interactions that govern activity in the environment. Microbe-microbe and microbe-environment interactions alter patterns of gene expression, activity, and growth, and so potential metabolism tells us little of the actual microbial processes that drive soil C-cycling at any particular place and time. The ‘realized activity’ of a microbe *in situ* depends on a complex pattern of biotic and abiotic interactions. Therefore, realized activity in the C-cycle might correlate only weakly with the metabolic potential of the community as inferred from phylogenetic conservation and metagenomic analyses (31, 32).

DNA-stable isotope probing (DNA-SIP) can assess the *in situ* activity of microorganisms within complex habitats based on patterns of C assimilation by identifying taxa that assimilate ^13^C from labeled substrates (33). DNA-SIP can track the assimilation of C from multiple substrates into thousands of microbial taxa in soil (34–36) with high specificity and sensitivity (36). We used multiple-window high-resolution DNA-SIP (MW-HR-SIP) to track ^13^C from nine distinct carbon sources through the soil bacterial food web over a period of 48 days (Fig. 1). These nine sources of C were selected because they varied in bioavailability and are common in soils. We initially defined bioavailability on the basis of solubility and hydrophobicity, which predicts sorption to particulate organic matter (37) and ability to transport across the cell membrane. The C sources we used were cellulose (a component of plant cell walls), glucose and xylose (monomers of cellulose and hemicellulose respectively), vanillin (a phenolic acid derived from lignin), glycerol and palmitic acid (components of plant lipids), an amino acid mixture (monomers of protein), and lactate and oxalate (major fermentation products from soil bacteria and fungi).

**Figure 1:**
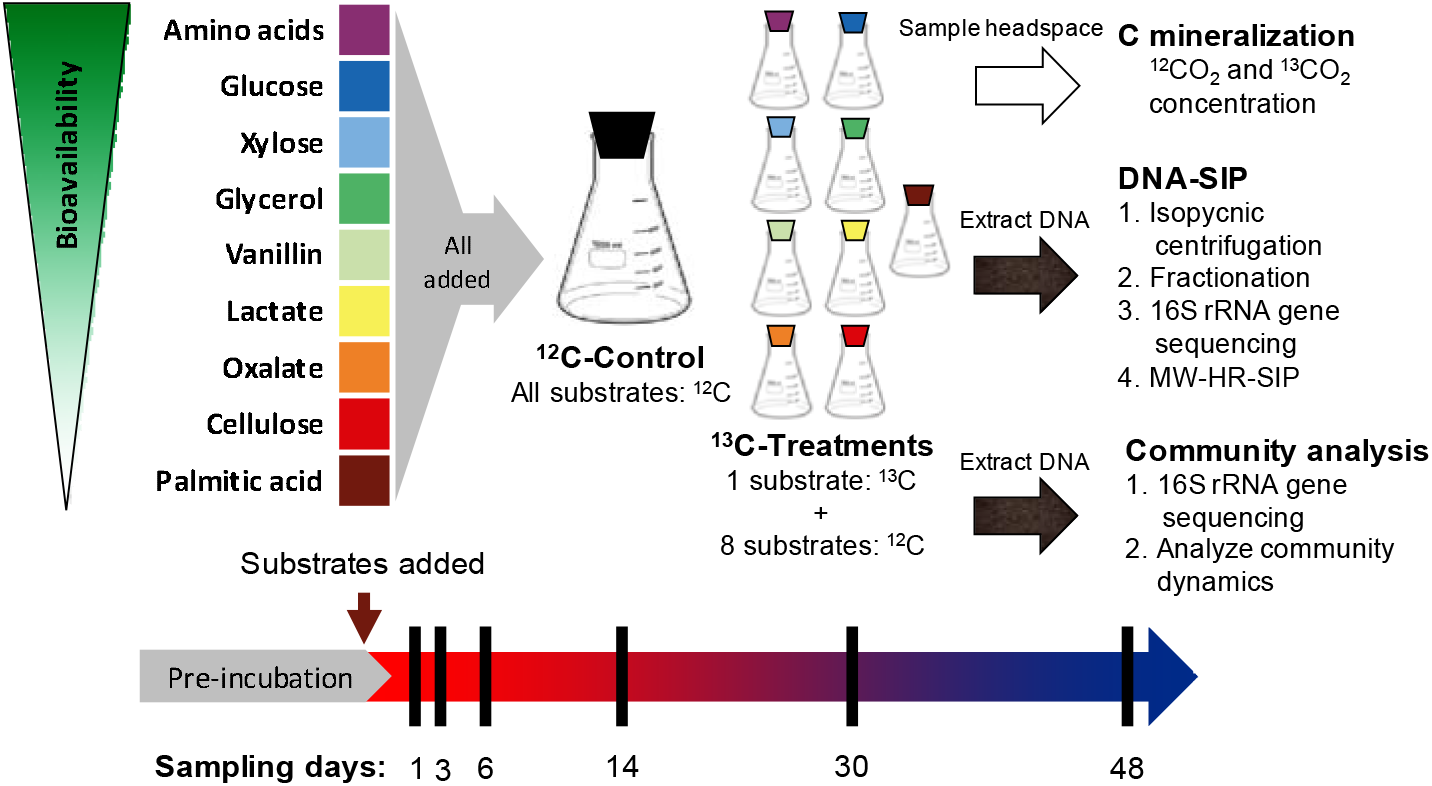
Diagram of the design of the DNA-SIP experiment. This experiment employed soil microcosms, each amended with all nine C sources (added at 0.4 mg C per gram of soil), only one of which was > 99% ^13^C-labeled in treatment microcosms. Control microcosms had all nine C sources added but none were isotopically labeled. Microcosms were destructively sampled at multiple timepoints (black bars) based on mineralization rates from preliminary experiment. Headspace samples were taken every 1-7 days. DNA extracted from microcosm soil was used both for DNA-SIP and whole bacterial community sequencing (unfractionated). Bioavailability estimated based on mineralization dynamics (Note that vanillin is highly hydrophobic and it occurred in both aqueous and nonaqueous (sorbed or solid) phases, but only the aqueous phase bioavailability is indicated here. Cellulose and palmitic acid were insoluble and were added as powders.).

Our multi-substrate design, where only the identity of the ^13^C-substrate was varied (Fig. 1), allowed us to systematically investigate C assimilation dynamics as a function of C source while maintaining identical conditions across samples. In addition, by sampling over time we focused on the ultimate fate of C within the bacterial community and not just the bacteria responsible for initial substrate breakdown. That is, our goal was not to differentiate primary assimilation from later stages, but rather to map bacterial guild structure by examining how C from diverse sources infiltrates the community over time. Guilds are groups of organisms that access resources in a similar manner (38). We used C assimilation dynamics to infer guild structure based on the source and timing of the ^13^C assimilated by bacterial taxa, such that bacteria within a guild exhibit similar patterns of C assimilation over time. We predicted that variations in C assimilation dynamics across bacterial taxa would be poorly explained by phylogenetic relatedness. Rather, ecological and physiological characteristics would be shared across taxa grouped into C assimilation guilds.

## Results and discussion

### Carbon source bioavailability explains carbon mineralization rates

We tracked C mineralization dynamics from each C source for up to 48 days by measuring ^13^CO_2_ production. Peak ^13^C mineralization rates occurred on day 1 for glucose and amino acids; day 2 for xylose, glycerol, and vanillin; day 3 for lactate; day 7 for oxalate; day 10 for cellulose; and day 14 for palmitic acid (Fig. 2A, Fig. S1). Cumulative ^13^C mineralization varied substantially between sources, being lowest for vanillin (42% C mineralized) and highest for oxalate (88% C mineralized, Fig. 2B, Fig. S2). As expected, total C mineralization (^13^C + ^12^C) was similar across all treatments (Fig. S1) since all microcosms received the same set of substrates with only the identity of the ^13^C-labeled substrate varied. These mineralization dynamics were confirmed in an independent experiment with soils from the same site (Fig. S2).

**Figure 2:**
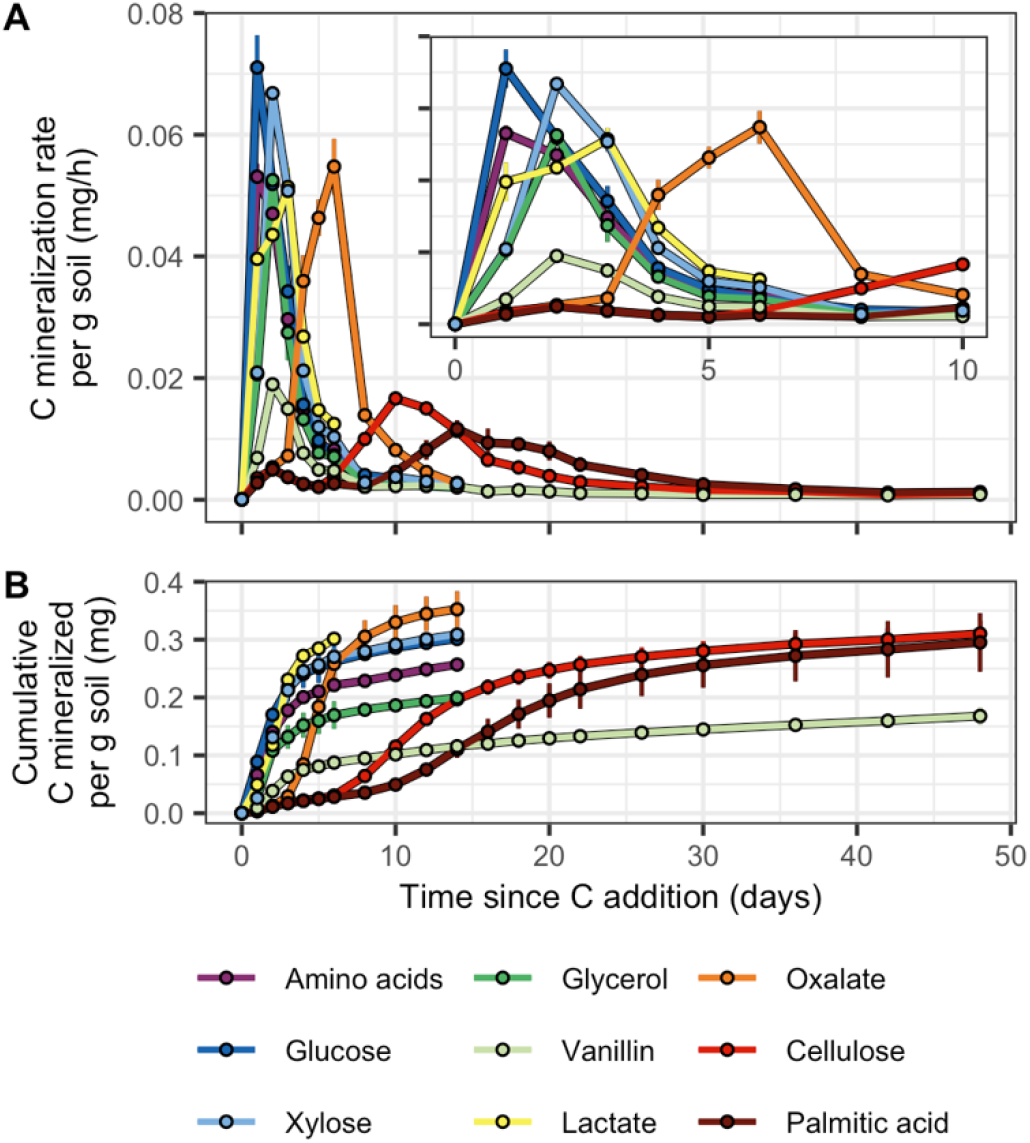
C source mineralization dynamics varied based on bioavailability. (A) ^13^C mineralization rates and (B) cumulative amounts of ^13^C mineralized per gram dry weight of soil. Inset for (A) displays a finer scale representation of the ^13^C mineralization rates over days 0-10. Error bars represent ± standard deviation among microcosm replicates (*n* = 3).

Substrate bioavailability predicts degradation rate in soil (39), and so we selected C sources that differ in their solubility and hydrophobicity (as defined by octanol-water partition coefficients predicted using the XLogP3 model (40)) as described in Supplemental Methods and indicated in Fig. 1. We found a significant and strong positive correlation between LogP and the day of peak C mineralization across C sources (Pearson’s r = 0.932, p-value = 0.001; Fig S3A) and a significant negative correlation between LogP and the maximum C mineralization rate (Pearson’s r = −0.855, p-value = 0.007; Fig S3B). These findings confirm that C sources with low bioavailability, such as cellulose (an insoluble polymer) and palmitic acid (a waxy fatty acid), are degraded slowly, while those with high bioavailability, such as glucose, xylose, amino acids, and lactate are degraded rapidly. Vanillin (an aromatic molecule) had intermediate bioavailability allowing it to be mineralized rapidly, however, its hydrophobicity favors adsorption to organic surfaces in soil (41, 42). Hence, while vanillin in the aqueous phase degrades rapidly, its sorption to organic matter may limit microbial access over time. Highly oxidized compounds such as oxalate favor allocation of C to catabolic processes, resulting in greater cumulative mineralization (43, 44). Given that LogP cannot be determined for an insoluble material such as cellulose, and that there was a strong relationship between LogP and day of peak mineralization (Fig. S3A), in subsequent analyses we defined bioavailability operationally based on observed day of peak mineralization.

### Non-cultivated bacteria are major players in soil C-cycling

The temporal dynamics of bacterial C assimilation were assessed by performing MW-HR-SIP for each of the nine ^13^C-labeled C sources over 48 days. Sampling times were selected based on the ^13^C mineralization dynamics for each substrate. A total of 12,394 unique bacterial OTUs were observed and 6,613 of these passed independent filtering on the basis of sparsity. From this set, MW-HR-SIP identified 1,286 “incorporators” (*i.e*., OTUs with significant evidence of ^13^C incorporation into DNA; Fig. 3; Supplemental Dataset). We also examined unfractionated DNA (*i.e*., DNA extracted directly from soil and not subjected to MW-HR-SIP) in order to evaluate overall change in microbial community composition over time. The majority of incorporators were also detected in unfractionated DNA (1,153, or about 90%), though 133 incorporators (~10%) were not observed in unfractionated DNA. This result indicates that rare OTUs, not readily detected in 16S rRNA gene surveys, are active participants in soil C-cycling. Density gradient centrifugation fractionates DNA by buoyant density with sequencing performed across many gradient fractions, and this approach translates into far greater sequencing depth than is typical in microbial community analyses, allowing for the detection of rare taxa not readily observed when sequencing unfractionated soil DNA.

**Figure 3:**
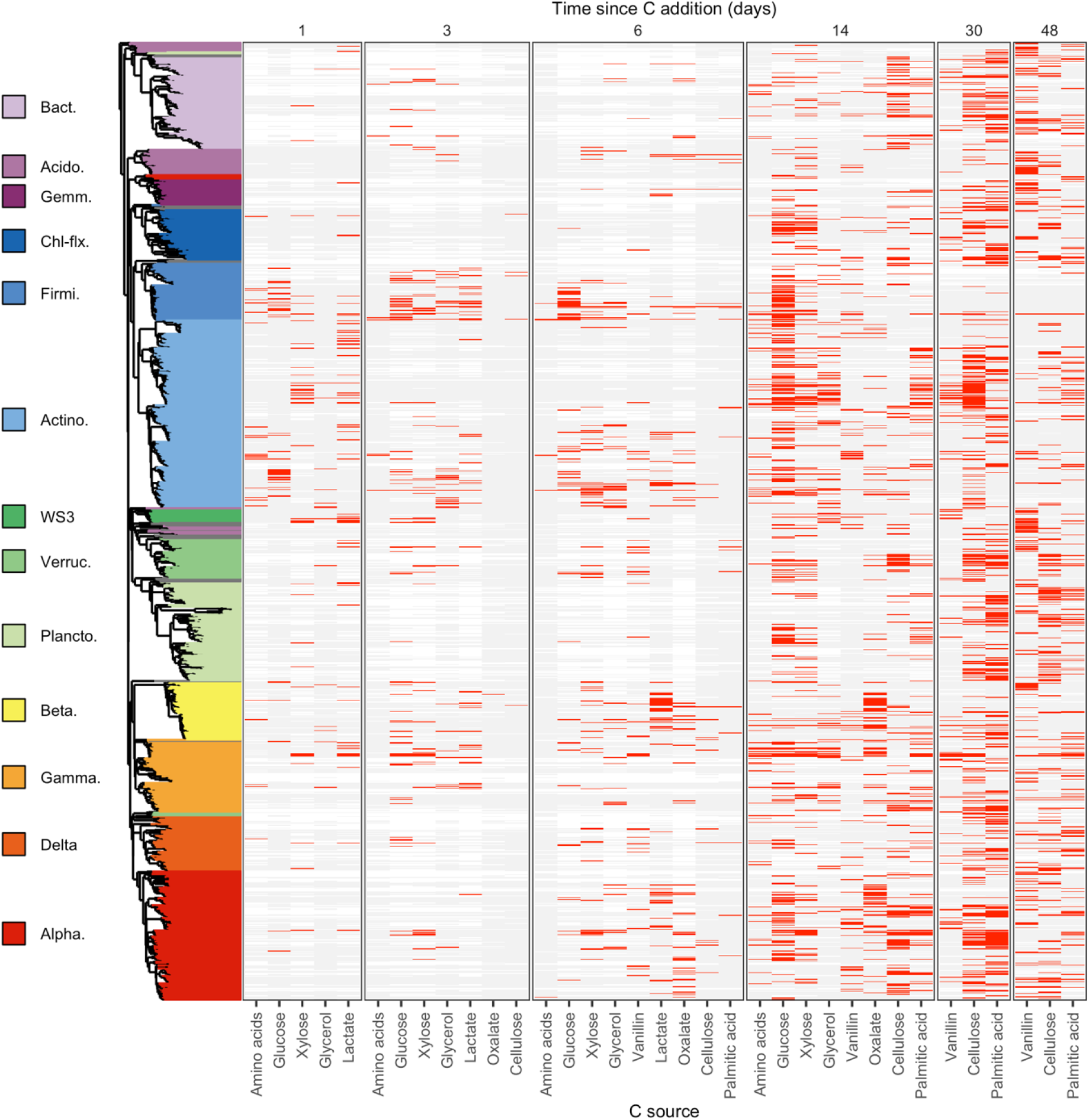
The dynamics of C assimilation varied across the 1,286 OTUs that exhibited significant ^13^C-labelling. OTUs are ordered by their phylogeny with Phylum, or Class for *Proteobacteria*, indicated by different colors in the phylogenetic tree. Only Phyla/Classes with 10 or more ^13^C-labeled OTUs are colored in this way with others colored grey. Each vertical block indicates ^13^C-labelling results for a single day as specified at the top. Each column within a block indicates results for a given substrate as specified on the bottom. Rows display the ^13^C-labelling dynamics for each OTU as follows: red bars indicate ^13^C-labelling, light grey bars indicate OTU detection in the gradient but no evidence of labeling, white indicates the OTU was not detected in the gradient. Phylum/Class abbreviations are Bact. = *Bacteroidetes*, Acido. = *Acidobacteria*, Gemm. = *Gemmatimonadetes*, Chl-flx. = *Chloroflexi*, Firmi. = *Firmicutes*, Actino. = *Actinobacteria*, Verruc. = *Verrucomicrobia*, Plancto. = *Planctomycetes*, Beta. = *Betaproteobacteria*, Gamma. = *Gammaproteobacteria*, Delta. = *Deltaproteobacteria*, Alpha. = *A lphaproteobacteria*.

Only 13.6% of the incorporators matched cultivated isolates (*i.e*., > 97% 16S rRNA gene sequence identity to known isolates), while 59.6% lacked a closely related isolate (*i.e*., < 97% 16S rRNA gene sequence identity), and 26.8% were only distantly related to any isolate (*i.e*., < 90% 16S rRNA gene sequence identity). C from low and intermediate bioavailability sources (*i.e*., cellulose, palmitic acid, and vanillin) was assimilated primarily by non-cultivated OTUs (Fig. S4). In contrast, C from high bioavailability sources (*i.e*., glucose, xylose, amino acids, glycerol, oxalate, and lactate) was assimilated primarily by OTUs matching known isolates (Fig. S4). Microbes that access C from low bioavailability sources are likely to have traits (*e.g*., slow growth, metabolic dependency, surface attachment) that diminish their tolerance for laboratory growth media and may limit their representation in culture collections (45, 46).

### Phylogenetic conservation of C assimilation dynamics

The ecophysiology of bacterial OTUs is often inferred from their phylogenetic relatedness (47–50). We therefore sought to determine if C assimilation activity could be predicted on the basis of phylogeny. We used two methods to test for the phylogenetic conservation in C assimilation dynamics. First, we calculated dissimilarity in assimilation dynamics between OTUs. Patterns of C assimilation were weakly conserved among closely related taxa and dissimilarity in C assimilation dynamics increased rapidly with phylogenetic distance (Fig. 4A-B). Even when analyzing C assimilation dynamics independently for each C source, phylogeny was a poor predictor of realized activity (Fig. S5). Second, we used consenTRAIT (49) to measure the phylogenetic depth (*i.e*., 16S rRNA gene evolutionary distance; τ_D_) at which patterns of C assimilation were conserved. We found τ_D_ values ranging from ~0.001 to 0.025 across the nine C sources (Fig. 4C), consistent with τ_D_ values of C substrate utilization measured by Martiny et al. (49). There was a significant correlation between τ_D_ and time of C assimilation (Pearson’s r = 0.791, p-value < 0.001; Fig. 4C). More specifically, phylogenetic depth was highest for C assimilation from vanillin, cellulose, and palmitic acid when assimilated at later time points, though the phylogenetic clustering of bacteria that assimilated C from these sources was not statistically significant across all time points (consenTRAIT; adjusted p-value ≥ 0.05; Fig. 4C). These results suggest that the assimilation of C from low bioavailability sources requires metabolic capabilities or traits that are more deeply conserved than those required to assimilate C from high bioavailability sources. The results also show that phylogenetic conservation of C-cycling process is highly variable, being higher for some processes than for others. Even within a single substrate such as cellulose, the degree of phylogenetic clustering depended on time of assimilation. High variation in phylogenetic clustering for taxa that access C from one source at different times suggests that phylogenetic conservation might be driven by shared ecology rather than shared catabolic pathways.

**Figure 4:**
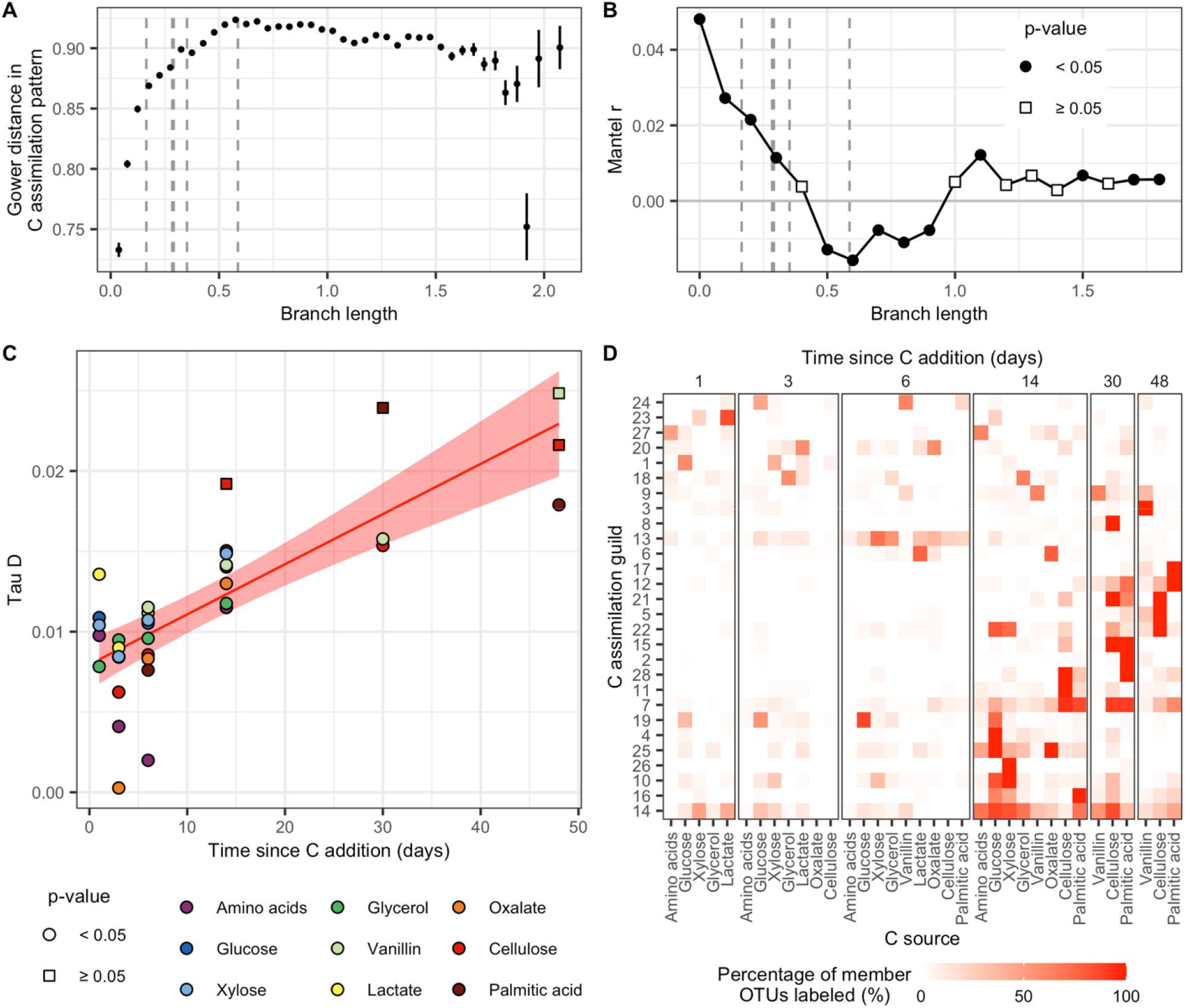
(A) The Gower’s distance in C assimilation dynamics (*i.e*., realized functional distance) between OTUs increases rapidly in relation to their phylogenetic distance. The vertical dashed lines indicate distance values that correspond to estimates of taxonomic ranks for genus, family, order, class, and phylum (from left to right). OTU pairs are grouped into phylogenetic distance bins with error bars indicating ± standard error. (B) Mantel test comparing realized functional distance and phylogenetic distance between OTUs with points colored by significance of correlation. Note very low Mantel r. (C) Phylogenetic depth (Tau D) of C assimilation from each substrate at each timepoint derived from consenTRAIT tests. All p-values are globally adjusted with the Benjamini-Hochberg correction. There is a positive correlation between Tau D and time (Pearson’s r = 0.79, p-value < 0.001). This linear relationship is represented by the red line with the ribbon representing the standard error. D) As phylogenetic relatedness poorly explained C assimilation dynamics, we instead grouped OTUs by their ^13^C-labeling pattern into guilds. Here C assimilation dynamics of the 28 guilds (ordered based on similarity in C assimilation dynamics) is displayed, with red color intensity indicating the percentage of member OTUs who have detectable ^13^C-labelling from each substrate on each day.

### C assimilation dynamics define bacterial guild structure

Since phylogenetic relatedness was a weak predictor of C assimilation dynamics in soil, we sought an alternative means to group bacteria into ecologically relevant clusters. We used unsupervised clustering to group incorporators into operationally defined guilds based on their ^13^C-labeling dynamics. This approach generated 28 guilds (Fig. 4D). Each guild contained between 15 and 122 incorporators and represented 3 to 13 different phyla (Supplemental Dataset). We then examined characteristic growth and ^13^C assimilation dynamics across guilds and related these characteristics with rRNA operon (*rrn*) copy number, a trait with known implications for bacterial ecology and C-cycling activity (51, 52). In all cases, characteristics and *rrn* copy number was averaged across all OTU within a guild (Supplemental Dataset).

After adjusting for multiple comparisons (*n* = 4), we found a positive correlation only between *rrn* copy number and the maximum log_2_ fold change in normalized abundance (max L_2_FC; Pearson’s r = 0.503, p-value = 0.025; Fig. 5D)). Max L_2_FC was defined as the maximum change in differential abundance before and after C sources were added to soil. High copy number guilds exhibited large changes in normalized abundance in response to C input, while low copy number guilds varied much less over time. In other words, guilds with high copy number exhibited highly dynamic populations while populations of guilds with low copy number were less dynamic. Since sequencing is compositional (53) and since several of the bacterial groups with the largest growth response had high copy number (Supplemental Dataset), relative abundance was normalized both by predicted *rrn* copy number and DNA yield for each sample. The *rrn* copy number of bacteria grown in culture correlates with maximal growth rate as well as C use efficiency, and has been proposed as a component of microbial life history strategies (51, 52). Our observation links the *rrn* copy number of guilds to their *in situ* population dynamics in response to C addition. There were two guilds with high *rrn* copy numbers (guilds 1 and 19), when removed from the analysis the relationship with maximum log_2_ fold change remained marginally significant (Pearson’s r = 0.458, p-value = 0.074; Fig. S6).

**Figure 5:**
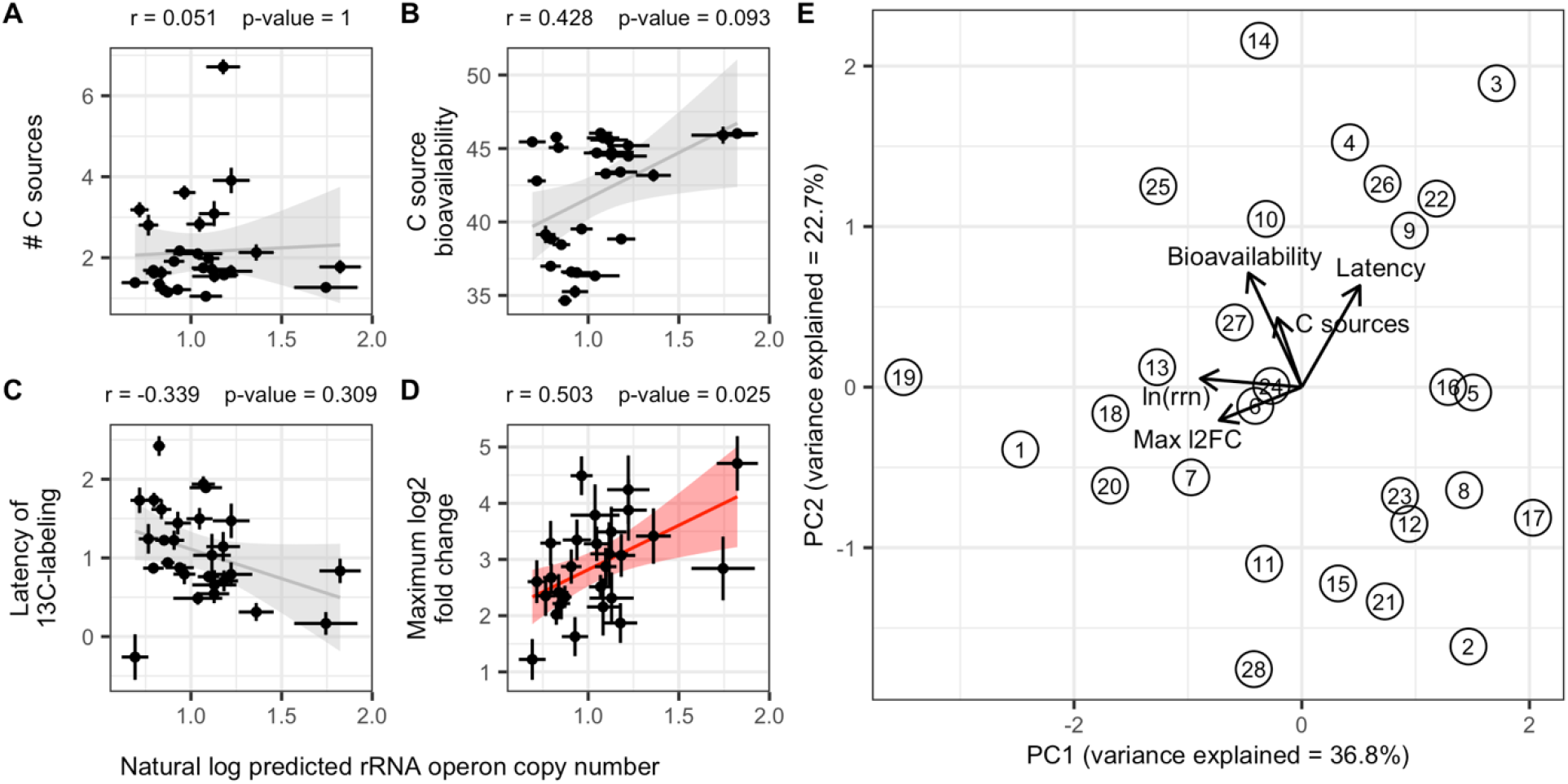
We propose that guild structure corresponds with life history strategies of bacteria. We tested whether the *rrn* copy number of guilds was correlated with other guild characteristics. A) The average number of C sources from which C was assimilated by each guild. B) The averaged bioavailability of the sources from which C was assimilated by each guild. Bioavailability is defined here as a unitless value determined operationally based on mineralization dynamics for each substrate, with higher values for more rapidly mineralized substrates. C) The averaged latency of C assimilation for each guild. Latency is defined operationally and it is proportional to the time delay between peak mineralization and the time of ^13^C assimilation into DNA for each C source. D) The dynamic growth response of each guild as measured by increase in abundance in response to substrate addition. Red and grey lines indicate the statistically significant and nonsignificant linear relationship between factors respectively with shading representing standard error. Pearson’s r and p-values for these relationships are listed above each plot. All p-values were corrected for multiple comparisons with Benjamini-Hochberg procedure (*n* = 4). Error bars for points indicate ± standard error across OTUs within each guild. E) The PCA differentiating guild structure on the basis of *rrn* copy number (ln(rrn)), C source bioavailability, latency of C assimilation, dynamic growth response (Max l2FC), and number of C sources (C sources). Circles represent guilds with numbers indicating identity and arrows representing loadings of each variable onto the principal component axes (PC1 and PC2).

We did not find any correlation between *rrn* copy number and the number of sources from which guilds derived their C (Pearson’s r = 0.051, p-value = 0.797; Fig. 5A), the averaged bioavailability of C source that led to ^13^C-labeling of the guilds (Pearson’s r = 0.428, p-value = 0.093; Fig 5B), or the ‘latency’ of ^13^C-labeling (Pearson’s r = −0.339, p-value = 0.077; Fig. 5C). Here, we defined bioavailability operationally as a function of the time when peak ^13^C-mineralization was observed, such that higher values are more available (Fig. S3). We defined latency as the relationship between the time of peak ^13^C-mineralization and the time ^13^C-labeling was first detected. A lower value for latency indicates that C assimilation took place when C mineralization was maximal, and hence the organism is likely using C directly from the source or is coupled tightly to the other organisms participating in mineralization. A higher value for latency indicates that C assimilation took place long after rates of C mineralization began to decline, making it more likely that the organism was assimilating C that had previously been transformed by microbial processing.

To further demonstrate the ecological relevance of this guild structure, we examined how phylogenetic distance among guild members, measured with the nearest taxon index (NTI), varied with respect to guild growth and ^13^C-assimilation dynamics. We expect that variation in selection pressure between guilds should alter phylogenetic clustering observed within guilds. For example, we predict greater competition for C from low bioavailability sources than from high bioavailable sources when C is added as a pulse. This expectation is based on the idea that consumption of low bioavailability substrates is constrained by diffusion and defined by spatial relationships. While high bioavailability pulsed resources (*e.g*., glucose) are available in great excess relative to the initial microbial biomass present in soil, causing competition to be minimal. Competitive exclusion is expected to cause phylogenetic overdispersion (low NTI). After adjusting for multiple comparisons (*n* = 4), we found positive correlations between NTI and both C source bioavailability (Pearson’s r = 0.64, p-value = 0.001; Fig. S7B) and population dynamics (Pearson’s r = 0.51, p-value = 0.022; Fig. S7D) of guilds. No correlation was observed with the number of C sources (Pearson’s r = 0.453, p-value = 0.062; Fig. S7A) nor C assimilation latency (Pearson’s r = −0.226, p-value = 0.994; Fig. S7C). These results indicate that competitive interactions within guilds are linked to the manner of C acquisition (i.e., the bioavailability of source C), and the growth dynamics of guild members.

While *rrn* copy number is one trait that can inform a guild’s ecology and relationship to SOC, traits often impose tradeoffs causing interactions that can confound simple correlations. We therefore used a principal component analysis (PCA) to explore variation in guild responses as a function of their characteristic growth and assimilation dynamics along with *rrn* copy number (Fig. 5E). The first principal component (PC1) explained 36.8% of variation in guild responses and this axis corresponds with variation in *rrn* copy number (43.1% loading), growth dynamics (max L_2_FC; 28.5% loading), latency (14.1% loading), C source bioavailability (11.9% loading), and number of C sources (2.48% loading). The second principal component (PC2) explains 22.7% of the variation in guild responses. This second axis largely corresponds with variation in C source bioavailability (44.5% loading), latency (35.2% loading), number of C sources (16.3% loading), growth dynamics (3.8% loading), and *rrn* copy number (0.3% loading). The overall guild structure suggests three extremes in guild characteristics, with certain characteristics maximized at each vertex (Fig. 5E). For example, guilds with the highest *rrn* copy number and growth dynamics (max L_2_FC) have low PC1 but intermediate PC2. Guilds assimilating C from the least bioavailable and fewest C sources have high PC1 and low PC2. Guilds with the highest latency in C assimilation, and greatest diversity of C assimilation patterns (C sources) have high PC1 and high PC2. These vertices represent the characteristic extremes of the bacterial guild structure, though many intermediate guilds exist.

We selected the most abundant OTUs (based on the summed normalized abundance across time) that represented the three vertices of the guild structure to illustrate the maximal differences between guilds. The most abundant OTU from guild 19 was OTU.7163, classified as a *Pseudomonas* species. This pseudomonad was estimated to have ~5 *rrn* operons, its abundance increased rapidly within a day following C input from a very low initial abundance, then dropped dramatically in abundance over time, and it assimilated C from glucose and cellulose at early time points (Fig. 6). Correspondingly, guild 19 has a high max L_2_FC, intermediate C source bioavailability, and low latency (Fig. 5). The most abundant OTU from guild 14 was OTU.22, classified as an *Agromyces* species. This microbacterium was estimated to have ~1 *rrn* operon, it increased in abundance after C input from a relatively high initial abundance, reaching maximal abundance after 6 days followed by a decline over time, and it was consistently ^13^C-labeled throughout the experiment assimilating C from multiple sources (Fig. 6). Correspondingly, guild 14 has an intermediate max L_2_FC, many C sources, and high latency (Fig. 5). The most abundant OTU from guild 2 was OTU.197, classified as a species of *Verrucomicrobia*. This verrucomicrobium was estimated to have ~2 *rrn* operons, it increased in abundance very slowly reaching maximal abundance after 30 days, remaining at relatively low abundance throughout, and assimilated C from glucose, vanillin, and cellulose at mostly later time points (Fig. 6). Correspondingly, guild 2 has a low max L_2_FC, low C source bioavailability, and low latency (Fig. 5). Generalized growth responses for all guilds are visualized in Figure S8.

**Figure 6:**
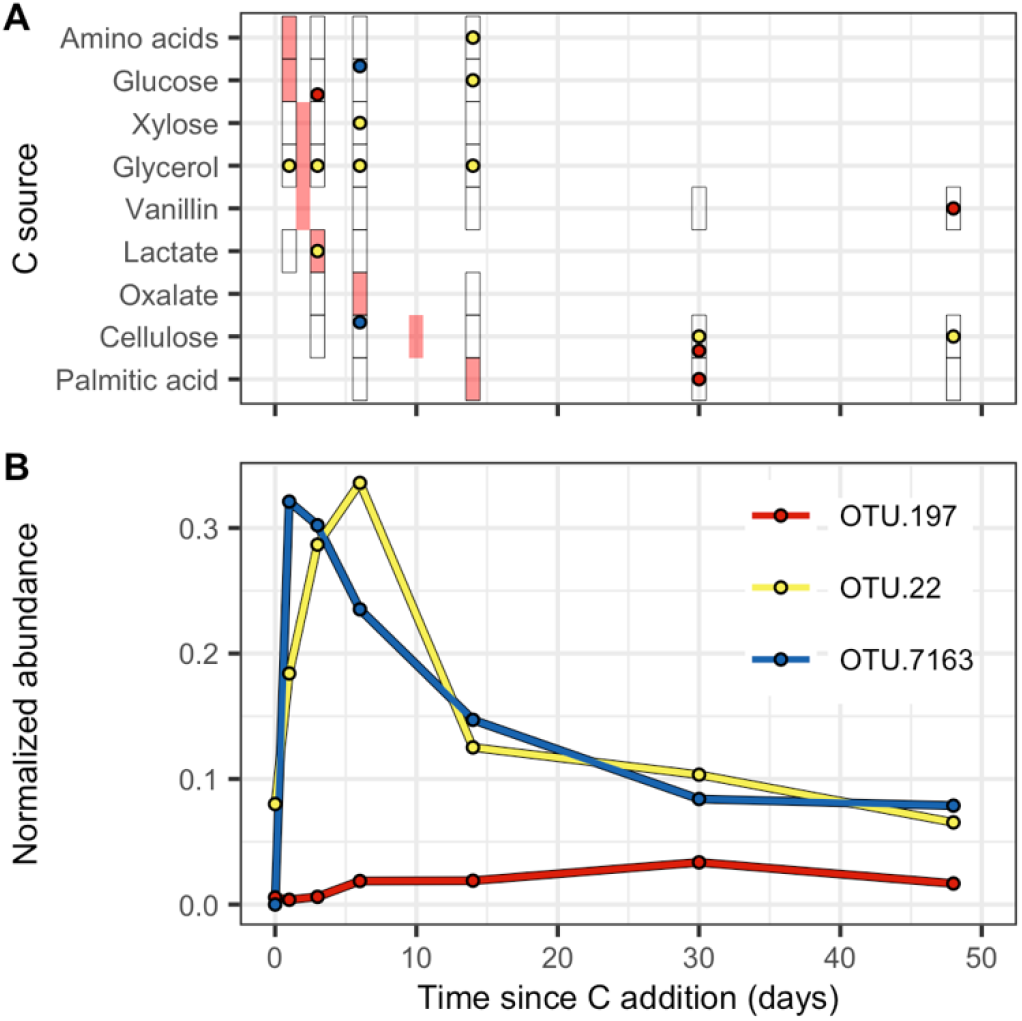
Growth and ^13^C-labeling dynamics of the most abundant OTUs from guilds at the extremes indicated on the PCA (Fig. 5E). A) The ^13^C-labeling pattern of the exemplar OTUs. Times when ^13^C-labeling was detected for each substrate are indicated by circles with colors indicating the different OTUs defined in panel B. Black rectangular borders indicate the timepoints in in which microcosms were harvested for DNA-SIP (*i.e*., where ^13^C-labeling can be detected). Red rectangles indicate the peak C mineralization timepoints for each C source (Fig. 2). B) The growth dynamics of the exemplar OTUs. Abundances are normalized by *rrn* copy number and DNA yield from the microcosms. OTU.7163 is a member of guild 19 and proposed to exemplify a ruderal strategist. OTU. 22 is a member of guild 14 and proposed to exemplify a competitor. OTU. 197 is a member of guild 2 and proposed to exemplify a stress tolerator.

It is clear from Figures 5, 6, and S8 that guilds differ in their growth response (max L_2_FC), and access to carbon (latency, C sources, bioavailability) and these differences suggest tradeoffs between growth dynamics and the ability to access C in the soil environment over time. Thus, the guild structure we identified can be interpreted in the context of life history theory, specifically Grime’s C-S-R framework (54). While Grime’s framework was conceived to describe plant life history strategies, it has recently been applied to microbial systems (55–57). Ruderal (R) strategists should have little investment in resource acquisition, being adapted for rapid growth in response to a sudden influx of nutrients. From our guild structure, ruderal bacteria are exemplified by the pseudomonads and bacilli of guild 19. These taxa have high *rrn* copy number, highly dynamic growth response (max L_2_FC), low latency, and C-labelling tends to occur immediately following nutrient addition, all together suggesting the ability to grow rapidly as a strategy for exploiting episodic nutrient pulses. Competitive (C) strategists are adapted for resource acquisition, likely allocating energy to features such as mycelia, biofilms, extracellular enzymes, siderophores, and antimicrobials that require substantial investment of energy but increase fitness under conditions defined by prolonged competition for resources.

Thus, competitors should grow more slowly (*i.e*., lower maximum growth rate), with less dynamic swings in population size. In our guild structure, competitors are exemplified by the *Agromyces* from guild 14. These taxa have higher latency suggesting the ability to access ^13^C following metabolic processing by other microbes, and they access C from a greater diversity of sources and over longer periods of time than ruderals. Both ruderals and competitors can be considered copiotrophs because of their ability to quickly respond to nutrient inputs (55), but these strategies would manifest differently in their population dynamics and sensitivity to nonequilibrium states (*e.g*., disturbance). Stress tolerators (S) were defined by Grime on the basis of abiotic stress (54), but we propose that for soil bacteria these organisms are adapted for life at low energy flux (*i.e*., conditions that would be stressful and result in starvation or dormancy for copiotrophs). In soil, we therefore propose that stress tolerators are synonymous with oligotrophs, organisms adapted to access low bioavailability C sources that would be otherwise unavailable to growth adapted copiotrophic organisms. Thus, in our guild structure, stress tolerant (S) strategists are those exemplified by the less well characterized *Verrucomicrobia* of guild 2. These taxa have low *rrn* copy number and very slow growth response, but also low latency and a tendency to access C from low bioavailability sources. While we focus on guilds 19, 14, and 2 to illustrate the presence of tradeoffs, Figure 5 illustrates that there is a wide diversity of strategies for accessing C from soil, including intermediate strategies not well binned strictly as ruderals, competitors, or stress tolerators (Fig. 5, S8). Similarly, members from the same taxonomic group may display dissimilar strategies. For example, while some *Pseudomonas* species exemplify the ruderal strategy, others can be found in guilds representing competitors or stress tolerators (Supplemental Dataset), demonstrating the physiological diversity and versatility of this genus.

Finally, we hope that is abundantly clear that the ecological tradeoffs that define fitness in soil are surely different from those that define fitness in other habitats (e.g. aquatic, gut, or extreme environments, etc.), and the categories we define here might not reflect well the determinants of microbial fitness in non-soil habitats. To illustrate this idea, consider the copiotroph-oligotroph framework, which was first used to describe the life history strategies of aquatic bacteria. Aquatic oligotrophs thrive at low substrate concentrations where substrate flux across the membrane is low and this imposes selection for high metabolic efficiency (58–65). Aquatic oligotrophic habitats are typically characterized by substrates that have low concentration at steady state (66). Under such equilibrium conditions, oligotroph growth rates are governed by substrate affinity as described by Michaelis-Menton kinetics (58). Soils, however, represent a highly dynamic, non-equilibrium system in which substrate availability varies over time and space in response to nutrient inputs, water availability, and substrate-mineral or substrate-organic matter interactions (67). Further, soil drying limits diffusion dramatically, resulting in periods of substrate limitation even if nutrients are otherwise abundant in bulk soil, while soil wetting enhances diffusive transport of soluble nutrients. Hence, wetting and drying of soil have major impacts on nutrient availability dynamics. Insoluble substrates, and those substrates adsorbed to surfaces, have low bioavailability being unable to cross the cytoplasmic membrane. Growth on these substrates is limited by the diffusion of enzymes and enzyme products between cells and their substrates. As a result, we predict that access to C from low bioavailability substrates favors oligotrophs in soil, because limitations on diffusion and transport should select for metabolic efficiency. Furthermore, diffusive fluxes in soil environments are favored by intimate contact between organism and substrate. Such close contacts persist in water films that coat soil aggregates, and remain in place as soils dry and diffusion is limited by increasing tortuosity. Hence, we predict that oligotrophic activity persists over a wide range of moisture availability in soils and that such activity is mediated by diverse collections of interacting organisms. Since aquatic oligotrophs are not constrained by diffusion and tortuosity, but rather by substrate concentration in solution, we expect they will be differentiated considerably from soil oligotrophs. In addition, we predict that access to C from more soluble, bioavailable substrates in soil favors copiotrophs with ruderal or competitive strategies. These substrates are ephemeral in nature outside of the rhizosphere, being transiently available in high concentration as a pulsed resource following wetting events. When soils are dry, water-soluble substrates are poorly available due to diffusional limitation. Wetting provides a flush that delivers these substrates into pore water where they are more accessible and where diffusion is less limited because tortuosity is low. The pulsed nature of soluble organic matter in soil pore water will favor rapid utilization of the C when nutrients are transiently present in excess, because the benefits of high affinity only matter when resources are present at low concentration and the benefits of high efficiency only matter when resources are present at steady state.

### High turnover of bacterial biomass

In terms of understanding microbial contributions to soil C cycling, much focus has been placed on primary microbial degradation of C derived from plants, and the degradation of plant biomass and exudates is typically considered in relation to microbial catabolism. However, recent evidence suggests that persistent SOM is largely composed from the products of microbial anabolic reactions (68). Hence, microbial turnover and mortality is likely of considerable importance in soil C-cycling. We observed rapid growth in response to C inputs, CO_2_ flux peaked at 2 days, and soil DNA increased over 6 days (a 52% increase), but there was also evidence of substantial turnover and mortality. For example, soil DNA declined considerably from day 6 through day 48 (a 20% decrease), and the normalized relative abundance of many taxa decreased over time. Of the ^13^C-labeled taxa, 387 were observed at high frequency over time (detection in 6 of 7 time points) in unfractionated soil DNA. Of these high frequency OTUs only 59 (15%) were observed to have a maximum in normalized relative abundance at the end of the experiment (*i.e*., no evidence of mortality), while 221 (57%) decreased by more than 50% in normalized relative abundance from an early peak in abundance. Decreases in normalized relative abundance were often dramatic and rapid (Fig. 6), particularly for guilds having high max L_2_FC (Fig. 5). In addition, among these frequently observed ^13^C-labeled taxa, a total of 202 (52%) were ^13^C-labeled at an early time point but were unlabeled by that same source at later time points. Loss of ^13^C-label can result from growth and isotope dilution over time, but when coupled to evidence of decreases in mineralization rate, decreases in DNA yield, and decreases in normalized relative abundance of most taxa over time, this result suggests mortality and turnover of ^13^C-DNA. Finally, the high latency for many ^13^C-labeled taxa (Fig. 5) indicates that many taxa became ^13^C-labeled indirectly by consuming microbial products rather than by direct assimilation of added C. Collectively these results argue that secondary processing, fueled by the products of microbial anabolic metabolism, comprise a major component of the C that cycles through soil communities. Based on our guild structure we would predict that much of this microbial biomass C ultimately flows through taxa having more competitive life history strategies (those with high latency in Fig. 5).

### Guild structure explains bacterial biogeography

To test for a linkage between guild structure and biogeography, we mapped incorporators to global bacterial surveys to determine whether guild membership explained variation in biogeography of soil bacteria. We employed RLQ analysis, a multivariate approach that finds correspondence between organism properties (R) and environmental parameters at various sites (Q; *e.g*., habitat or soil pH) based on organismal abundance within sites (L) (69, 70). We mapped incorporator OTUs and their guild membership to two 16S rRNA gene surveys of soil bacterial diversity spanning continental (QIITA study 619) and global scales (QIITA study 928) (71). Guild membership explained significant variation in microbial relative abundance across environmental parameters (Table S1; Fig. S9). For simplicity, guilds were divided into three groups based on C assimilation profile. Group D assimilated C primarily from dissolved substrates of high bioavailability (glucose, xylose, amino acids, glycerol, lactate, and oxalate). Group P assimilated C primarily from particulate substrates of low bioavailability (cellulose and palmitic acid). Group V assimilated C from vanillin; this group was treated separately as it was likely present in both aqueous and noaqueous forms with transition between these two states changing over time (owing to its insolubility and tendency to sorb to SOM). Guilds were further grouped by time of labelling: early (E) and late (L). At continental scales, guilds that primarily used dissolved substrates at late timepoints were associated with forest soils, and soils having high organic C and N (DL; Fig. S9A). At global scales, these guilds were positively associated with latitude (DL; Fig. S9B), likely as a result of the northern concentration of forest ecosystems and SOM at global scales. The guilds most strongly associated with tropical and subtropical broadleaf forests were those that assimilated C from particulate substrates early (PE; Fig. S9B). The guilds that were most strongly associated with temperate grasslands, savannas, and shrubland were those that assimilated C from diverse substrates (DL, PL, and VL; Fig. S9B). Guilds that used C from particulate substrates and vanillin at later timepoints were most closely associated with desert biomes (PL and VL; Fig. S9A). These results show that patterns of microbial C assimilation can be linked to patterns of microbial biogeography and that guild structure has relevance across large biogeographical scales.

## Conclusions

We conducted a multi-substrate DNA-SIP experiment to measure the *in situ* activities of bacteria in the soil C-cycle. We show that degradation dynamics and patterns of bacterial C assimilation depended on the bioavailability of C inputs. In total, 1286 bacteria were found to incorporate C derived from 9 different sources, and most of these bacteria were distantly related to cultivated isolates. We found that C assimilation dynamics in soil had low phylogenetic conservation among bacteria. Instead, C assimilation dynamics allowed us to group bacteria into guilds whose properties were generally consistent with well-known life history strategies. We also show that this guild structure can explain variation in bacterial biogeography. Our results demonstrate that life history strategies are likely linked to C assimilation dynamics. SOM is largely derived from the products of anabolic microbial metabolism (14), and therefore microbial assimilation and turnover of C is a primary determinant of C fate in soil. Based on our results, we hypothesize that C fate in soil is not determined solely, or even primarily, by the metabolic pathways that enable its catalysis, but rather by the life history traits that govern microbial growth and C acquisition *in situ*. As a result, life history theory could be useful in predicting the C-cycle activity of microbial communities (18). While more research is necessary to better understand the genetic and ecological underpinnings of life history strategies in soil bacteria, our results add to the growing body of knowledge indicating the importance of life history theory to bacterially mediated terrestrial C-cycling.

## Materials and methods

### Soil microcosm experiments

Soil microcosms were designed and incubated with ^13^C-labeled and unlabeled substrates as previously described (35) except that C amendment consisted of nine substrates: cellulose, xylose, glucose, glycerol, vanillin, palmitic acid, amino acid mixture, lactate, and oxalate (SI; Table S2). For most substrates, octanol-water partition coefficient (LogP) was determined using XLogP3 through PubChem (40). Since amino acids were added as a mixture, this substrate LogP was averaged across the 20 common amino acids. LogP is not available for cellulose as it is insoluble. Each substrate was added at 0.4 mg C g^-1^ soil. Only one of the nine substrates was ^13^C-labeled in each treatment while the remainder were unlabeled (Fig. 1). Soil was from Penn Yan, New York, USA, with site and collection methods previously described (35, 72) and outlined in SI. ^13^C and ^12^C mineralization dynamics were determined based on concentrations of ^12^CO_2_ and ^13^CO_2_ in microcosm headspace measured over time as detailed in SI. Soil from microcosms were destructively sampled depending on substrate mineralization dynamics over days 1, 3, 6, 14, 30, and 48 after substrate addition.

### DNA extraction and isopycnic centrifugation

DNA extraction and isopycnic centrifugation was conducted as previously described (35) and detailed in SI. In short, for each treatment, size selected DNA (≥ 4 kb) (73) was centrifuged in a CsCl density gradient and 100 μl fractions were collected. An aliquot of DNA taken before isopycnic centrifugation was used for whole microcosm bacterial community sequencing (*i.e*., unfractionated DNA).

### 16S rRNA gene amplification and sequencing

We amplified and sequenced the V4 region of the 16S rRNA gene as previously described (74) and detailed in SI using dual indexed primers (515f and 806r) developed by Kozich *et al*. (75). 16S rRNA amplicon libraries were processed as described previously (35) and detailed in SI. OTUs were clustered at 97% sequence similarity. Raw sequencing reads can be accessed at the NCBI Short Read Archive (accession PRJNA668741).

### Identifying incorporators

OTUs that incorporated ^13^C into their DNA (*i.e*., incorporators) were identified with MW-HR-SIP (36) using development code for the HTSSIP R package (76). In short, we used DESeq2 (77) to identify OTUs significantly enriched in the “heavy” gradient fractions of the ^13^C-labeled treatments compared to the “heavy” gradient fractions in corresponding ^12^C-control. Incorporators were defined as OTUs with a log_2_ fold enrichment greater than 0.25 and a Benjamini-Hochberg adjusted p-value less than 0.1. We used the overlapping BD windows 1.70-1.73, 1.72-1.75, and 1.74-1.77 g/ml, which were shown via simulations to have the highest sensitivity across all tested scenarios while not losing specificity (36). Independent sparsity filtering was conducted for each BD window to minimize the number of comparisons performed.

The taxonomic novelty of incorporators was determined by aligning OTU representative sequences to the Silva all-species living tree project small subunit database version 123, representing all sequenced type strains curated at that time (78). Alignment was performed using nucleotide BLAST (79) with an expected value cutoff of 1E-20. Incorporators with no hits ≥ 97% sequence identity were considered to have no closely related isolates, while those with no hits ≥ 90% were considered to have no related isolates. Statistical analyses were conducted in R version 3.1.2 (80).

### Analyses of phylogenetic conservation

Fasttree v2.1.9 (81) was used to infer a maximum likelihood phylogeny from the 16S rRNA sequence alignment of representative OTUs. The phylogeny was rooted with *Sulfolobus solfataricus* DSM1616 (Genbank accession X90478) as the outgroup. Phylogenetic conservation of realized function was examined both by comparing functional distance with phylogenetic distance and testing for a phylogenetic signal with a modified version of consenTRAIT (49) (https://github.com/nick-youngblut/consenTRAIT).

### Analysis of guild structure

Incorporators were grouped into guilds based on their ^13^C-labelling patterns as determined by log_2_ fold change in ^13^C-labeled treatment fractions versus corresponding unlabeled control fractions. Specifically, clustering was performed by: *i*) calculating pairwise Gower distances for each OTU pair, *ii*) hierarchically clustering (UPGMA agglomeration method) based on the resulting distance matrix, *iii*) generating a dendrogram of the hierarchical associations, *iv*) detecting clusters in the dendrogram using the *cutreeHybrid* function in the DynamicTreeCut R package (82).

The *rrn* copy number of each OTU was predicted as described previously (83). Briefly, pplacer v1.1 (84) was used to insert OTU representative sequences into a 16S rRNA gene reference phylogeny of bacteria with known *rrn* copy number. The resulting phylogeny was used by guppy v1.1 to predict *rrn* copy numbers of each OTU based on its relatedness to the reference taxa. The natural log of the copy number was used for all comparisons (51).

Four guild C assimilation and growth characteristics were calculated. All characteristics were first calculated for individual OTUs then averaged within each guild. The number of C sources from which C was assimilated was simply the number of C sources from which an OTU was ^13^C-labeled at any timepoint. Since logP is not available for cellulose, C source bioavailability was defined operationally based on the day of maximal ^13^C mineralization rate for each substrate (as detailed in SI). The C source bioavailability was determined as the average of all sources from which ^13^C was assimilated. Latency of C assimilation was determined by the natural log of the ratio between the first day of ^13^C-labelling and the day of peak C mineralization for each C source. This value was then averaged across all sources from which C was assimilated. The maximum log_2_ fold change in OTU abundance in the unfractionated DNA was calculated following abundance normalization to minimize bias due to compositional data. Normalization included two steps: normalizing for predicted *rrn* copy number and normalizing by sample DNA yield. OTU relative abundances were first divided by their predicted rRNA operon copy number, then by the estimate copy number for the entire community. Within each timepoint, normalized relative abundances were then multiplied by the average DNA yield across the replicate microcosms. The extracted DNA yield (ng DNA g^-1^ dry weight soil) was quantified with the Quant-iT PicoGreen dsDNA Assay Kit (Thermo Fisher Scientific, Waltham, MA, USA). We recognize that this calculation does not take into account inflation in DNA yield due to the growth of fungi and other organisms not accounted for in our study, however we believe that these effects will be small compared to the effects of bacterial growth, particularly as we are simply comparing between bacterial OTUs within this study. Untreated bulk soils were used as our baseline abundance, with OTUs undetected in these soils assigned the lowest abundance measured. For each OTU, the log_2_ fold change in abundance between this baseline and the timepoint when abundance was highest was then calculated. If abundance at all timepoints was less than the baseline or undetected, the maximum abundance was assigned the baseline abundance, making the log_2_ fold change = 0 for such OTUs. If an OTU is ^13^C-labeled, it is growing on the substrates provided. A decrease in abundance at a time when taxa are labeled indicates either that the rate of mortality for the population exceeds the division rate (deaths > births), that labelling occurred prior to growth decline, or that normalization was not entirely successful at eliminating all variance due to compositional sequencing.

After averaging across all member OTUs within each guild, we measured the relationship between the natural log of the rRNA operon copy number and each of the four C assimilation characteristics using the Pearson’s correlation. We also measured the relationship between all four C assimilation characteristics and the nearest taxon index (NTI) within guilds. NTI was calculated using R package picante (85). *rrn* copy number and all four of characteristics were used in a principal component analysis (PCA) to more fully assess the variation in these characteristics across guilds.

### Mapping incorporators to biogeography datasets

Detailed descriptions of the independent datasets and their analysis for this study are found in SI. In short, OTU count tables, representative 16S rRNA gene sequences, and metadata was downloaded from https://qiita.ucsd.edu/ for QIITA studies 619 (continental dataset) and 928 (global dataset). Incorporators were mapped to external dataset OTUs separately for each study with the mothur alignment tool (86) using the Silva reference alignment as a template. To examine the biogeography of guilds at continental (QIITA 619) and global (QIITA 928) scales, guilds were grouped by assimilation dynamics (as described in results). Then for each QIITA dataset, guild designations (R) of mapped OTUs were used in combination with their OTU count tables (L) and sample metadata (Q) for RLQ and forth corner analyses (70, 87) as detailed in SI.

## Acknowledgements

We thank Chuck Pepe-Ranney, Ashley Campbell, Marquessa Henry, and Braulio Castillo, Anay Hindupur, and Jillian Waters for their help with experimental design and implementation.

## References

1. E. G. Jobbágy, R. B. Jackson, The vertical distribution of soil organic carbon and its relation to climate and vegetation. Ecol. Appl. 10, 423–436 (2000).

2. V. Brovkin, et al., Effect of anthropogenic land-use and land-cover changes on climate and land carbon storage in CMIP5 projections for the twenty-first century. J. Climate 26, 6859–6881 (2013).

3. J. P. W. Scharlemann, E. V. J. Tanner, R. Hiederer, V. Kapos, Global soil carbon: understanding and managing the largest terrestrial carbon pool. Carbon Manag. 5, 81–91 (2014).

4. Z. Luo, G. Wang, E. Wang, Global subsoil organic carbon turnover times dominantly controlled by soil properties rather than climate. Nat. Commun. 10, 3688 (2019).

5. M. A. Bradford, et al., Managing uncertainty in soil carbon feedbacks to climate change. Nat. Clim. Change 6, 751–758 (2016).

6. R. Lal, Soil carbon sequestration impacts on global climate change and food security. Science 304, 1623–1627 (2004).

7. R. Lal, Soil health and carbon management. Food Enrgy. Security 5, 212–222 (2016).

8. D. W. Reeves, The role of soil organic matter in maintaining soil quality in continuous cropping systems. Soil Till. Res. 43, 131–167 (1997).

9. P. Smith, Soils as carbon sinks: the global context. Soil Use Manage. 20, 212–218 (2004).

10. M. Heimann, M. Reichstein, Terrestrial ecosystem carbon dynamics and climate feedbacks. Nature 451, 289–292 (2008).

11. K. E. O. Todd-Brown, et al., Changes in soil organic carbon storage predicted by Earth system models during the 21st century. Biogeosciences 11, 2341–2356 (2014).

12. Y. Luo, T. F. Keenan, M. Smith, Predictability of the terrestrial carbon cycle. Glob. Change Biol. 21, 1737–1751 (2015).

13. W. R. Wieder, et al., Carbon cycle confidence and uncertainty: Exploring variation among soil biogeochemical models. Glob. Change Biol. 24, 1563–1579 (2018).

14. J. Schimel, S. Schaeffer, Microbial control over carbon cycling in soil. Front. Microbiol. 3, 348 (2012).

15. K. E. O. Todd-Brown, et al., Causes of variation in soil carbon simulations from CMIP5 Earth system models and comparison with observations. Biogeosciences 10, 1717–1736 (2013).

16. W. R. Wieder, et al., Explicitly representing soil microbial processes in Earth system models. Global Biogeochem. Cy. 29, 1782–1800 (2015).

17. G. Wang, W. M. Post, M. A. Mayes, Development of microbial-enzyme-mediated decomposition model parameters through steady-state and dynamic analyses. Ecol. Appl. 23, 255–272 (2013).

18. W. R. Wieder, A. S. Grandy, C. M. Kallenbach, G. B. Bonan, Integrating microbial physiology and physio-chemical principles in soils with the MIcrobial-MIneral Carbon Stabilization (MIMICS) model. Biogeosciences 11, 3899–3917 (2014).

19. B. N. Sulman, R. P. Phillips, A. C. Oishi, E. Shevliakova, S. W. Pacala, Microbe-driven turnover offsets mineral-mediated storage of soil carbon under elevated CO2. Nat. Clim. Change 4, 1099–1102 (2014).

20. W. R. Wieder, G. B. Bonan, S. D. Allison, Global soil carbon projections are improved by modelling microbial processes. Nat. Clim. Change 3, 909–912 (2013).

21. L. A. Hug, et al., A new view of the tree of life. Nat. Microbiol. 1, 16048 (2016).

22. K. G. Lloyd, A. D. Steen, J. Ladau, J. Yin, L. Crosby, Phylogenetically novel uncultured microbial cells dominate earth microbiomes. mSystems 3, e00055–18 (2018).

23. A. D. Steen, et al., High proportions of bacteria and archaea across most biomes remain uncultured. ISME J. 13, 3126–3130 (2019).

24. M. G. I. Langille, et al., Predictive functional profiling of microbial communities using 16S rRNA marker gene sequences. Nat. Biotechnol. 31, 814–821 (2013).

25. K. P. Aßhauer, B. Wemheuer, R. Daniel, P. Meinicke, Tax4Fun: predicting functional profiles from metagenomic 16S rRNA data. Bioinformatics 31, 2882–2884 (2015).

26. M. Brbić, et al., The landscape of microbial phenotypic traits and associated genes. Nucleic Acids Res. 44, 10074–10090 (2016).

27. J. B. H. Martiny, S. E. Jones, J. T. Lennon, A. C. Martiny, Microbiomes in light of traits: A phylogenetic perspective. Science 350, aac9323 (2015).

28. J. Choi, et al., Strategies to improve reference databases for soil microbiomes. ISME J. 11, 829–834 (2017).

29. N. Fierer, et al., Cross-biome metagenomic analyses of soil microbial communities and their functional attributes. P. Natl. Acad. Sci. U.S.A. 109, 21390–21395 (2012).

30. J. Alneberg, et al., Genomes from uncultivated prokaryotes: a comparison of metagenome-assembled and single-amplified genomes. Microbiome 6, 173 (2018).

31. D. L. Wohl, S. Arora, J. R. Gladstone, Functional redundancy supports biodiversity and ecosystem function in a closed and constant environment. Ecology 85, 1534–1540 (2004).

32. R. Evans, et al., Defining the functional traits that drive bacterial decomposer community productivity. ISME J. 11, 1680–1687 (2017).

33. M. G. Dumont, J. C. Murrell, Stable isotope probing — linking microbial identity to function. Nat. Rev. Microbiol. 3, 499–504 (2005).

34. B. A. Hungate, et al., Quantitative microbial ecology through stable isotope probing. Appl. Environ. Microb. 81, 7570–7581 (2015).

35. C. Pepe-Ranney, A. N. Campbell, C. N. Koechli, S. Berthrong, D. H. Buckley, Unearthing the ecology of soil microorganisms using a high resolution DNA-SIP approach to explore cellulose and xylose metabolism in soil. Front. Microbiol. 7, 703 (2016).

36. N. D. Youngblut, S. E. Barnett, D. H. Buckley, SIPSim: A modeling toolkit to predict accuracy and aid design of DNA-SIP experiments. Front. Microbiol. 9, 570 (2018).

37. D. Sijm, R. Kraaij, A. Belfroid, Bioavailability in soil or sediment: exposure of different organisms and approaches to study it. Environ. Pollut. 108, 113–119 (2000).

38. D. Simberloff, T. Dayan, The guild concept and the structure of ecological communities. Annu. Rev. Ecol. Syst. 22, 115–143 (1991).

39. B. Berg, K. Hannus, T. Popoff, O. Theander, Changes in organic chemical components of needle litter during decomposition. Long-term decomposition in a Scots pine forest. Can. J. Botany 60, 1310–1319 (1982).

40. T. Cheng, et al., Computation of octanol-water partition coefficients by guiding an additive model with knowledge. J. Chem. Inf. Model. 47, 2140–2148 (2007).

41. T. Polubesova, Y. Chen, R. Navon, B. Chefetz, Interactions of hydrophobic fractions of dissolved organic matter with Fe^3+^ - and Cu^2+^-montmorillonite. Environ. Sci. Technol. 42, 4797–4803 (2008).

42. M. Keiluweit, M. Kleber, Molecular-level interactions in soils and sediments: the role of aromatic π-systems. Environ. Sci. Technol. 43, 3421–3429 (2009).

43. J. B. Brant, E. W. Sulzman, D. D. Myrold, Microbial community utilization of added carbon substrates in response to long-term carbon input manipulation. Soil Biol. Biochem. 38, 2219–2232 (2006).

44. S. Manzoni, P. Taylor, A. Richter, A. Porporato, G. I. Ågren, Environmental and stoichiometric controls on microbial carbon-use efficiency in soils. New Phytol. 196, 79–91 (2012).

45. K. Alain, J. Querellou, Cultivating the uncultured: limits, advances and future challenges. Extremophiles 13, 583–594 (2009).

46. V. H. T. Pham, J. Kim, Cultivation of unculturable soil bacteria. Trends Biotechnol. 30, 475–484 (2012).

47. J. T. Lennon, Z. T. Aanderud, B. K. Lehmkuhl, D. R. Schoolmaster Jr., Mapping the niche space of soil microorganisms using taxonomy and traits. Ecology 93, 1867–1879 (2012).

48. S. A. Placella, E. L. Brodie, M. K. Firestone, Rainfall-induced carbon dioxide pulses result from sequential resuscitation of phylogenetically clustered microbial groups. P. Natl. Acad. Sci. U.S.A. 109, 10931–10936 (2012).

49. A. C. Martiny, K. Treseder, G. Pusch, Phylogenetic conservatism of functional traits in microorganisms. ISME J. 7, 830–838 (2013).

50. K. L. Dolan, J. Peña, S. D. Allison, J. B. H. Martiny, Phylogenetic conservation of substrate use specialization in leaf litter bacteria. PLOS ONE 12, e0174472 (2017).

51. B. R. Roller, T. M. Schmidt, The physiology and ecological implications of efficient growth. ISME J. 9, 1481–1487 (2015).

52. N. Fierer, M. A. Bradford, R. B. Jackson, Toward an ecological classification of soil bacteria. Ecology 88, 1354–1364 (2007).

53. G. B. Gloor, J. M. Macklaim, V. Pawlowsky-Glahn, J. J. Egozcue, Microbiome datasets are compositional: and this is not optional. Front. Microbiol. 8, 2224 (2017).

54. J. P. Grime, Evidence for the existence of three primary strategies in plants and its relevance to ecological and evolutionary theory. Am. Nat. 111, 1169–1194 (1977).

55. J. I. Prosser, et al., The role of ecological theory in microbial ecology. Nat. Rev. Microbiol. 5, 384–392 (2007).

56. A. Ho, et al., Conceptualizing functional traits and ecological characteristics of methane-oxidizing bacteria as life strategies. Env. Microbiol. Rep. 5, 335–345 (2013).

57. N. Fierer, Embracing the unknown: disentangling the complexities of the soil microbiome. Nat. Rev. Microbiol. 15, 579–590 (2017).

58. D. K. Button, Biochemical basis for whole-cell uptake kinetics: specific affinity, oligotrophic capacity, and the meaning of the Michaelis constant. Appl. Environ. Microb. 57, 2033–2038 (1991).

59. D. K. Button, Nutrient-limited microbial growth kinetics: overview and recent advances. A. Van Leeuw. J. Microb. 63, 225–235 (1993).

60. C. E. Zobell, C. W. Grant, Bacterial utilization of low concentrations of organic matter. J. Bacteriol. 45, 555–564 (1943).

61. J. R. Postgate, J. R. Hunter, The survival of starved bacteria. Microbiology 29, 233–263 (1962).

62. H. W. Jannasch, Growth of marine bacteria at limiting concentrations of organic carbon in seawater. Limnol. Oceanogr. 12, 264–271 (1967).

63. S. I. Kuznetsov, G. A. Dubinina, N. A. Lapteva, Biology of oligotrophic bacteria. Annu. Rev. Microbiol. 33, 377–387 (1979).

64. Y. Akagi, U. Simidu, N. Taga, Growth responses of oligotrophic and heterotrophic marine bacteria in various substrate concentrations, and taxonomic studies on them. Can. J. Microbiol. 26, 800–806 (1980).

65. D. K. Button, Nutrient uptake by microorganisms according to kinetic parameters from theory as related to cytoarchitecture. Microbiol. Mol. Biol. R. 62, 636–645 (1998).

66. C. Arnosti, Microbial extracellular enzymes and the marine carbon cycle. Annu. Rev. Mar. Sci. 3, 401–425 (2011).

67. P. Sollins, P. Homann, B. A. Caldwell, Stabilization and destabilization of soil organic matter: mechanisms and controls. Geoderma 74, 65–105 (1996).

68. C. Liang, W. Amelung, J. Lehmann, M. Kästner, Quantitative assessment of microbial necromass contribution to soil organic matter. Glob. Change Biol. 25, 3578–3590 (2019).

69. S. Dolédec, D. Chessel, C. J. F. ter Braak, S. Champely, Matching species traits to environmental variables: a new three-table ordination method. Environ. Ecol. Stat. 3, 143–166 (1996).

70. S. Dray, et al., Combining the fourth-corner and the RLQ methods for assessing trait responses to environmental variation. Ecology 95, 14–21 (2014).

71. S. T. Bates, et al., Examining the global distribution of dominant archaeal populations in soil. ISME J. 5, 908–917 (2011).

72. S. T. Berthrong, D. H. Buckley, L. E. Drinkwater, Agricultural management and labile carbon additions affect soil microbial community structure and interact with carbon and nitrogen cycling. Microb. Ecol. 66, 158–170 (2013).

73. N. D. Youngblut, D. H. Buckley, Intra-genomic variation in G + C content and its implications for DNA stable isotope probing. Env. Microbiol. Rep. 6, 767–775 (2014).

74. S. E. Barnett, N. D. Youngblut, D. H. Buckley, Soil characteristics and land-use drive bacterial community assembly patterns. FEMS Microbiol. Ecol. 96, fiz194 (2020).

75. J. J. Kozich, S. L. Westcott, N. T. Baxter, S. K. Highlander, P. D. Schloss, Development of a dual-index sequencing strategy and curation pipeline for analyzing amplicon sequence data on the MiSeq Illumina sequencing platform. Appl. Environ. Microb. 79, 5112–5120 (2013).

76. N. D. Youngblut, S. E. Barnett, D. H. Buckley, HTSSIP: An R package for analysis of high throughput sequencing data from nucleic acid stable isotope probing (SIP) experiments. PLOS ONE 13, e0189616 (2018).

77. M. I. Love, W. Huber, S. Anders, Moderated estimation of fold change and dispersion for RNA-seq data with DESeq2. Genome Biol. 15, 550 (2014).

78. P. Yarza, et al., The All-Species Living Tree project: A 16S rRNA-based phylogenetic tree of all sequenced type strains. Syst. Appl. Microbiol. 31, 241–250 (2008).

79. S. F. Altschul, W. Gish, W. Miller, E. W. Myers, D. J. Lipman, Basic local alignment search tool. J. Mol. Biol. 215, 403–410 (1990).

80. R Core Team, R: A language and environment for statistical computing (2018).

81. M. N. Price, P. S. Dehal, A. P. Arkin, FastTree: Computing large minimum evolution trees with profiles instead of a distance matrix. Mol. Biol. Evol. 26, 1641–1650 (2009).

82. P. Langfelder, B. Zhang, S. Horvath, Defining clusters from a hierarchical cluster tree: the Dynamic Tree Cut package for R. Bioinformatics 24, 719–720 (2008).

83. S. W. Kembel, M. Wu, J. A. Eisen, J. L. Green, Incorporating 16S gene copy number information improves estimates of microbial diversity and abundance. PLOS Comput. Biol. 8, e1002743 (2012).

84. F. A. Matsen, R. B. Kodner, E. V. Armbrust, pplacer: linear time maximum-likelihood and Bayesian phylogenetic placement of sequences onto a fixed reference tree. BMC Bioinformatics 11, 538 (2010).

85. S. W. Kembel, et al., Picante: R tools for integrating phylogenies and ecology. Bioinformatics 26, 1463–1464 (2010).

86. P. D. Schloss, et al., Introducing mothur: open-source, platform-independent, community-supported software for describing and comparing microbial communities. Appl. Environ. Microb. 75, 7537–7541 (2009).

87. S. Dray, D. Chessel, J. Thioulouse, Co-inertia analysis and the linking of ecological data tables. Ecology 84, 3078–3089 (2003).

