## Supplemental Materials and Methods for "Multi-substrate DNA stable isotope probing reveals guild structure in bacterially mediated soil carbon cycling"

*Soil microcosm experiments*

Soils were collected from an organic farm in Penn Yan, New York, USA. Detailed description of the field site can be found in Berthrong *et al.* 2013 (1) and Pepe-Ranney *et al.* 2016 (2). Soil used to make microcosms for the preliminary substrate mineralization experiment were collected on August 26^th^, 2014, while soil used for the DNA-SIP microcosms were collected on October 27^th^, 2014. The field consisted of 6 strips with an alternating crop rotation scheme. At each strip, 15 soil cores (5 cm deep) were collected spaced about 1 m apart along a North-South transect. Soil cores were sieved to 2 mm, homogenized together, then stored at 4˚C overnight. An aliquot for bulk soil bacterial community sequencing was stored at -80˚C. Soil moisture was measured gravimetrically (2).

Soil incubations with ^13^C-labeled and unlabeled substrates were performed as previously described (2) except that C amendment consisted of nine substrates: cellulose, xylose, glucose, glycerol, vanillin, palmitic acid, amino acid, lactate, and oxalate. These nine substrates were chosen as they vary in their bioavailability initially determined by solubility and hydrophobicity. Hydrophobicity was defined by the octanol-water partition coefficients predicted using the XLogP3 model (3) and reported in PubChem. It is impossible to determine logP for cellulose, since it is insoluble. We therefore removed cellulose from the comparison of C mineralization dynamics and bioavailability (Fig. S3). As the amino acids were added as a homogenous mixture, we used an average of the predicted LogP of all 20 common amino acids.

Isotopically labeled substrates were acquired from various sources (Table S2) but were all >99% ^13^C-labeled. In order to maintain consistent metabolic dynamics between treatments, all substrates were added to each microcosm, but each treatment different in which particular substrate was ^13^C-labeled with the other 8 unlabeled (Fig. 1). A set of ^12^C-control microcosms, consisting of all substrates added but unlabeled, were created for each sampling day. Additionally, a set of H_2_O-control microcosms were created which did not receive any substrates. Microcosms consisted of 15 g dry weight soil in a 250 mL sterile glass Erlenmeyer flask sealed with a sterile rubber stopper. To each microcosm (except H_2_O-controls), each substrate was added such that 0.4 mg of C was added per g dry weight of soil. This mass of carbon has been shown to be sufficient for sensitivity of HR-SIP methodologies (2). Cellulose and palmitic acid, which are insoluble in water, were autoclaved sterilized and then evenly dispersed across the soil surface after passing through a 250 µm sieve. All other substrates were dissolved in a 2.9% Murashige Skoog basal salt mixture (Sigma Aldrich M5524) at a volume to bring water moisture to 50% then evenly spread over soil surface with a sterile Mucosal Atomization Device (Mountainside Medical Equipment, Marcy, NY, USA). Three replicate microcosms were created for each sampling day. Microcosms were stored at room temperature in the dark until sampling. Microcosms were sampled at multiple timepoints based on their carbon mineralization dynamics in the preliminary study (Fig. S2). ^13^C-glucose, ^13^C-xylose, ^13^C-amino acids, and ^13^C-glycerol microcosms were sampled 1, 3, 6 and 14 days after substrate addition. ^13^C-lactate microcosms were sampled on days 1, 3, and 6. ^13^C-oxalate was sampled on days 3, 6, and 14. ^13^C-vanillin microcosms were sampled on days 6, 14, 30 and 48. ^13^C-cellulose microcosms were sampled on days 3, 6, 14, 30, and 48. ^13^C-palmitic acid microcosms were sampled on days 6, 14, 30, and 48. ^12^C-control microcosms were sampled on all days and H_2_O-control microcosms sampled on day 48. Sampling was destructive, so separate microcosms were created for each sampling. Soils were removed from the microcosms using a sterile spatula and stored immediately at -80˚C. Whenever possible, microcosms were stored and handled in a randomized order so as to minimize batch effects. Microcosm headspace was collected repeatedly throughout the experiment from the set of replicate microcosms from each treatment to be sampled last. 250 µl of headspace was collected and run on a GCMS-QP2010S (Shimadzu, Kyoto, Japan) containing a Carboxen 1010 PLOT column (Supelco, Bellefonte, PA, USA) with helium as the carrier gas. All microcosms headspaces were then flushed with air through a 0.22 µm filter to limit oxygen depletion.

*DNA extraction and isopycnic centrifugation*

DNA extraction and isopycnic centrifugation was conducted as previously described (2). DNA was extracted from 0.25 g of frozen soil using a modified Griffiths protocol (4). DNA was quantified with a Quant-iT PicoGreen dsDNA Assay Kit (Thermo Fisher Scientific, Waltham, MA, USA). Replicate DNA extractions were performed for each treatment in order to get over 5 µg of DNA (usually about 5 replicate extractions). An aliquot of DNA extract was used directly for whole bacterial community sequencing (later referred to as “unfractionated”) but was first purified with illustra MicroSpin G-50 Columns.

Prior to isopycnic centrifugation, DNA was size-selected (≥ 4 kb) with a BluePippin with 0.75% agarose, dye free, low range cassettes (Sage Science, Beverly, MA, USA). Size selection was done in order to reduce the centrifugation time required for all DNA fragments to reach sedimentation equilibrium (5). Density gradients consisted of 5 µg size-selected DNA in 1X TE buffer, 430 µl gradient solution (1.69 g ml^-1^ CsCl, 15 mM Tris-HCL (pH 8.0), 15 mM KCl, and 15 mM EDTA) with enough extra 1X TE buffer to bring total volume to 450 µl. Gradients were generated in 4.7 ml OptiSeal ultracentrifuge tubes (#361621, Beckman Coulter, Brea, CA, USA). Ultracentrifugation was conducted at 55000 RPM at 20˚C for ≥ 66 hr in a TLA-110 rotor with an Optima MAX-E ultracentrifuge (Beckman Coulter, Brea, CA, USA). Gradient fractions of ~100 µl were collected by pumping water into the top of the gradient with displaced fractions collected from a pierced hole in the tube bottom. Fractions were desalted with the Agencourt Ampure XP system (Beckman Coulter, Brea, CA, USA).

*16S rRNA gene amplification and sequencing*

We amplified the V4 region of the 16S rRNA gene as previously described (6) using dual indexed primers (515f and 806r) developed by Kozich *et al.* (7). Polymerase chain reaction (PCR) amplification was performed in 25 µl triplicate reactions with 2 µl template (0-5 ng DNA), 13.1 µl Q5 hot start high fidelity master mix (New England Biolabs, Ipswich, MA, USA) mixed 1:0.025 v/v with 4X Quant-iT PicoGreen reagent (Thermo Fisher Scientific, Waltham, MA, USA), 2.5 µl mixed 10X primers, and 7.4 µl PCR grade water. PCR conditions were 95˚C for 2 min, followed by 30 cycles of 95˚C for 20 sec, 55˚C for 15 sec, and 72˚C for 10 sec, and followed by 72˚C for 5 min. Triplicate successful PCR reactions were pooled and normalized with the Invitrogen SequalPrep Normalization Plate Kit (Thermo Fisher Scientific, Watham, MA, USA). Pooled amplicon libraries (up to 192 samples each) were gel-purified with the Wizard SV Gel and PCR Clean-up System (Promega, Madison, WI, USA) and sequenced on the Illumina MiSeq platform with paired end 2x250 bp V2 kit at the Cornell Biotechnology Resource Center (Ithaca, NY, USA). Nine libraries were sequenced in total. Raw sequencing reads can be accessed at the NCBI Short Read Archive (accession PRJNA668741).

Separately for each sequencing library, forward and reverse sequences were merged using PEAR (8) and demultiplexed using a custom script. Reads were filtered using alignment based quality filtering (SILVA SEED database, maximum homopolymer length 8, and maximum expected error 1) with mothur (9). Reads classified as mitochondria, chloroplasts, or *Archaea* were removed. At this point all libraries were combined. OTUs were clustered to 97% sequence identity and chimeras were removed using USEARCH (10). OTU taxonomy was assigned based on SILVA release 111 with the uclust algorithm through QIIME (11).

*Guild C assimilation and growth characteristics*

Four guild C assimilation and growth characteristics were calculated. All characteristics were first calculated for individual OTUs then averaged across OTUs within each guild. The number of C sources from which C was assimilated was simply the number of C sources from which an OTU was ^13^C-labeled at any timepoint. Since logP is not available for cellulose, C source bioavailability was defined operationally based on the day of maximal ^13^C mineralization rate for each substrate. This value was calculated as 48 (length of the experiment in days) minus the day of maximal ^13^C mineralization rate for each substrate. Thus, earlier mineralized C sources had higher bioavailability. For each OTU, bioavailability was averaged across all sources from which ^13^C was assimilated. Latency of C assimilation was calculated by taking the natural log of the ratio between the first day of ^13^C-labelling and the day of peak C mineralization for each C source for each OTU. This value was then averaged across all sources from which C was assimilated. The maximum log_2_ fold change in OTU abundance in the unfractionated DNA was calculated following abundance normalization to minimize bias due to compositional data. Normalization included two steps: normalizing for predicted *rrn* copy number and normalizing by sample DNA yield. OTU relative abundances were first divided by their predicted rRNA operon copy number, then by the estimate copy number for the entire community. Within each timepoint, normalized relative abundances were then multiplied by the average DNA yield across the replicate microcosms. The extracted DNA yield (ng DNA g^-1^ dry weight soil) was quantified with the Quant-iT PicoGreen dsDNA Assay Kit (Thermo Fisher Scientific, Waltham, MA, USA). Untreated bulk soils were used as our baseline abundance, with OTUs undetected in these soils assigned the lowest abundance measured. For each OTU, the log_2_ fold change in abundance between this baseline and the timepoint when abundance was highest was then calculated. If abundance at all timepoints was less than the baseline or undetected, the maximum abundance was assigned the baseline abundance, making the log_2_ fold change = 0 for such OTUs. If an OTU is ^13^C-labeled, it is growing on the substrates provided. A decrease in abundance at a time when taxa are labeled indicates either that the rate of mortality for the population exceeds the division rate (deaths > births), that labelling occurred prior to growth decline, or that normalization was not entirely successful at eliminating all variance due to compositional sequencing.

*Mapping incorporators to independent datasets to assess biogeography*

OTU count tables, representative 16S rRNA gene sequences, and metadata for studies 619 (continental dataset) and 928 (global dataset) were downloaded from QIITA (https://qiita.ucsd.edu/). Incorporator OTUs were mapped to these datasets with the mothur alignment tool (9) using the Silva reference alignment as a template. OTUs mapped at 97% sequence identity were retained. For simplicity, guilds were grouped based on the form of the substrates from which they assimilated C (D = dissolved, including glucose, xylose, amino acids, glycerol, oxalate, and lactate; V = vanillin; and P = particulate, including cellulose and palmitic acid) and time of C assimilation (E = early, first half of timepoints; L = late, second half of timepoints). For each dataset, guilds designations (R) of mapped OTUs were used in combination with their OTU count tables (L) and sample metadata (Q) for RLQ and forth corner analyses (12, 13). For the RLQ analysis, the correspondence analysis was applied to the L table, and a PCA was applied to the R and Q tables. The environmental variables in the PCA of the Q table were weighted by site coefficients from the correspondence analysis of the L table. Monte-Carlo permutations (n = 9999) were used to test for a trait-environment relationship, with the neutral model set as type 6 (12). The type 6 null model was also used for the fourth corner analysis (n = 9999) and the Benjamini-Hochberg correction for multiple hypotheses was applied. Both RLQ and fourth corner analyses were conducted with the ade4 R package (14). In order to reduce the number of multiple hypotheses in the fourth corner analysis, only guilds with an absolute cumulative loading > 0.4 for principal components 1 and 2 were included in both fourth corner and RLQ analyses. Testing for significant bivariate relationships between guilds and environmental parameters with the fourth corner analysis was not fruitful due to the high number of multiple hypotheses and possibly because the associations were not bivariate (data not shown).

**References**

1.	S. T. Berthrong, D. H. Buckley, L. E. Drinkwater, Agricultural management and labile carbon additions affect soil microbial community structure and interact with carbon and nitrogen cycling. *Microb. Ecol.* **66**, 158–170 (2013).

2.	C. Pepe-Ranney, A. N. Campbell, C. N. Koechli, S. Berthrong, D. H. Buckley, Unearthing the ecology of soil microorganisms using a high resolution DNA-SIP approach to explore cellulose and xylose metabolism in soil. *Front. Microbiol*. **7**, 03 (2016).

3.	T. Cheng, *et al.*, Computation of octanol−water partition coefficients by guiding an additive model with knowledge. *J. Chem. Inf. Model.* **47**, 2140–2148 (2007).

4.	R. I. Griffiths, A. S. Whiteley, A. G. O’Donnell, M. J. Bailey, Rapid method for coextraction of DNA and RNA from natural environments for analysis of ribosomal DNA- and rRNA-based microbial community composition. *Appl. Environ. Microbiol*. **66**, 5488–5491 (2000).

5.	N. D. Youngblut, D. H. Buckley, Intra-genomic variation in G + C content and its implications for DNA stable isotope probing. *Environ. Microbiol. Rep.* **6**, 767–775 (2014).

6.	S. E. Barnett, N. D. Youngblut, D. H. Buckley, Soil characteristics and land-use drive bacterial community assembly patterns. *FEMS Microbiol. Ecol.* **96**, fiz194 (2019).

7.	J. J. Kozich, S. L. Westcott, N. T. Baxter, S. K. Highlander, P. D. Schloss, Development of a dual-index sequencing strategy and curation pipeline for analyzing amplicon sequence data on the MiSeq Illumina sequencing platform. *Appl. Environ. Microbiol*. **79**, 5112–5120 (2013).

8.	J. Zhang, K. Kobert, T. Flouri, A. Stamatakis, PEAR: a fast and accurate Illumina Paired-End reAd mergeR. *Bioinformatics* **30**, 614–620 (2014).

9.	P. D. Schloss, *et al.,* Introducing mothur: open-source, platform-independent, community-supported software for describing and comparing microbial communities. *Appl. Environ. Microbiol.* **75**, 7537–7541 (2009).

10.	R. C. Edgar, Search and clustering orders of magnitude faster than BLAST. *Bioinformatics* **26**, 2460–2461 (2010).

11.	J. G. Caporaso, *et al.,* QIIME allows analysis of high-throughput community sequencing data. *Nat. Methods* **7**, 335–336 (2010).

12.	S. Dray, P. Choler, S. Dolédec, P. R. Peres-Neto, W. Thuiller, S. Pavoine, C. J. F. ter Braak, Combining the fourth-corner and the RLQ methods for assessing trait responses to environmental variation. *Ecology* **95**, 14–21 (2014).

13.	S. Dray, D. Chessel, J. Thioulouse, Co-inertia analysis and the linking of ecological data tables. *Ecology* **84**, 3078–3089 (2003).

14.	S. Dray, A. B. Dufour, The ade4 package: implimenting the duality diagram for ecologists. *J. Stat. Softw.* **22**, 1–20 (2007).
