## Supplemental Tables and Figures for "Multi-substrate DNA stable isotope probing reveals guild structure in bacterially mediated soil carbon cycling"

**Table S1:** p-values for global significance permutation tests of RLQ associations. Model 2 tests for a link between traits and abundances (assuming an association between abundance and environmental variables). Model 4 tests for a link between abundances and environmental variables (assuming an association between traits and abundance). Significance for both tests indicates that traits, abundances, and environmental parameters are associated. For each test, 9999 permutation replicates were conducted.

| **Survey scale** | **QIITA study ID** | **pvalue** | |
| --- | --- | --- | --- |
|  |  | **Model 2** | **Model 4** |
| Continental | 619 | 7e-4 | 1e-4 |
| Global | 928 | 1e-4 | 7e-4 |

**Table S2:** Isotopically labeled substrates used in the DNA-SIP experiment.

| **Substrate** | **Isotopic labeling** | **^13^C enrichment** | **Manufacturer** | **Manufacturer Location** |
| --- | --- | --- | --- | --- |
| cellulose | * | * | * | * |
| D-xylose | ^13^C_5_ | 99% | Omicron | South Bend, IN, USA |
| D-glucose | ^13^C_6_ | 99% | Cambridge Isotopes | Tewksbury, MA, USA |
| glycerol | ^13^C_3_ | 99% | Sigma Isotec | Miamisburg, OH, USA |
| vanillin | ring-^13^C_6_ | 99% | Sigma Isotec | Miamisburg, OH, USA |
| palmitic acid | ^13^C_16_ | 99% | Sigma Isotec | Miamisburg, OH, USA |
| algal amino acid mixture | ^13^C | 98% | Cambridge Isotopes | Tewksbury, MA, USA |
| lactate | ^13^C_3_ | 99% | Sigma Isotec | Miamisburg, OH, USA |
| oxalate | ^13^C_2_ | 99% | Sigma Isotec | Miamisburg, OH, USA |

* Bacterial cellulose was produced by *Gluconoacetobacter xylinus* as described in Pepe-Ranney *et al.,* (2016)

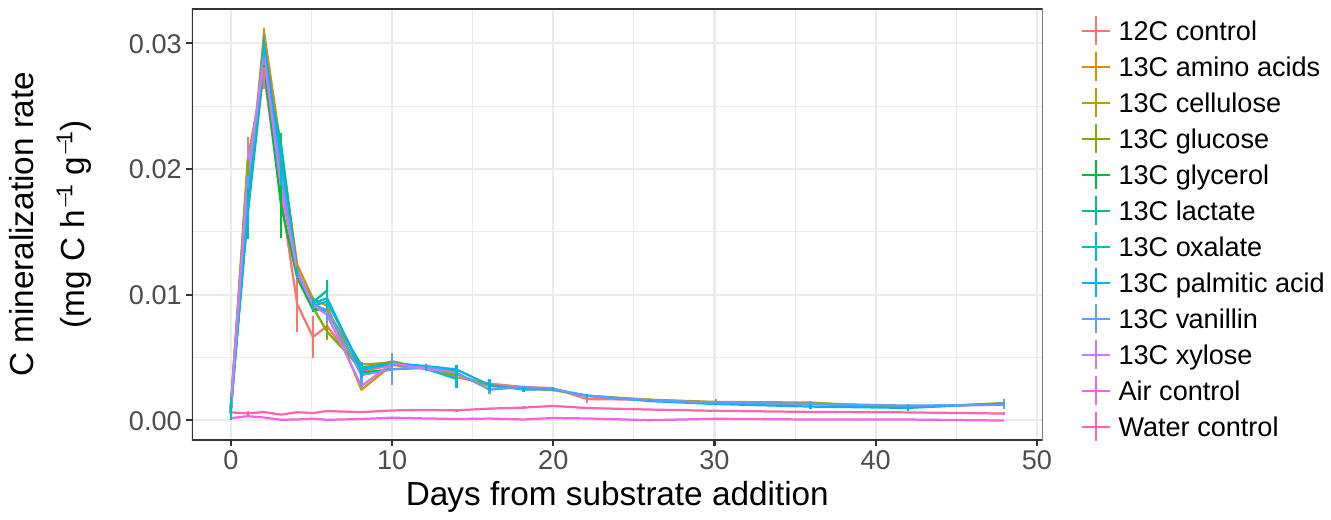

**Figure S1:** Total C (summed ^12^C and ^13^C) was consistent across all microcosms. This result is expected as all treatments are identical and the only variable is the identity of the ^13^C-labeled substrate. Error bars represent ± standard deviation among microcosm replicates (*n* = 3).

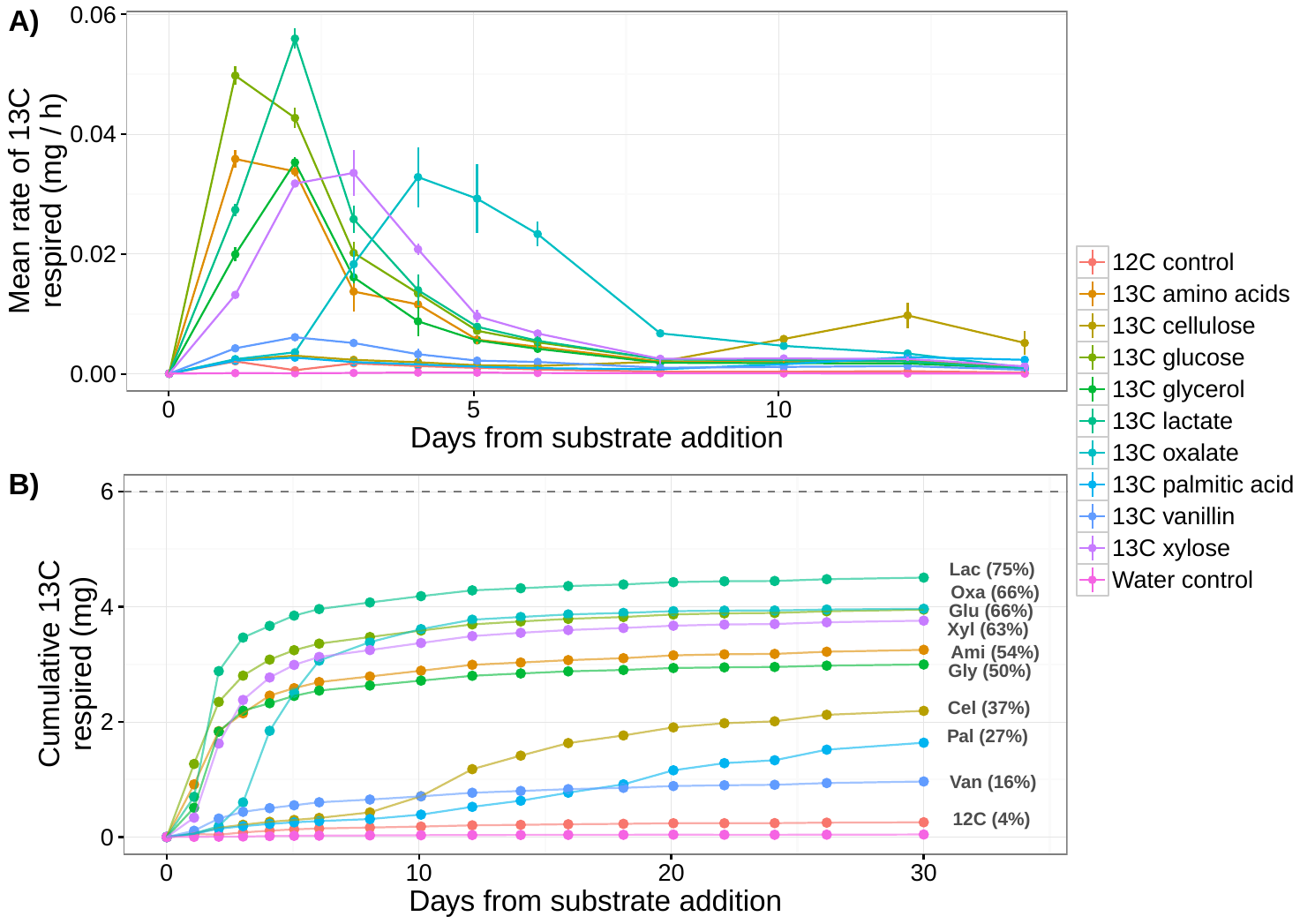

**Figure S2:** Substrate ^13^C mineralization dynamics for pilot experiment using the same soils as the main study with A) mineralization rate and B) cumulative mineralization over time. For clarity, only days 0-14 are shown for mineralization rate. Error bars indicate ± standard deviation among microcosm replicates (*n* = 4). The dashed line in B) represents the total amount of ^13^C added to each microcosm (6 mg) and the percentages indicate the percentage of the ^13^C mineralized by day 30.

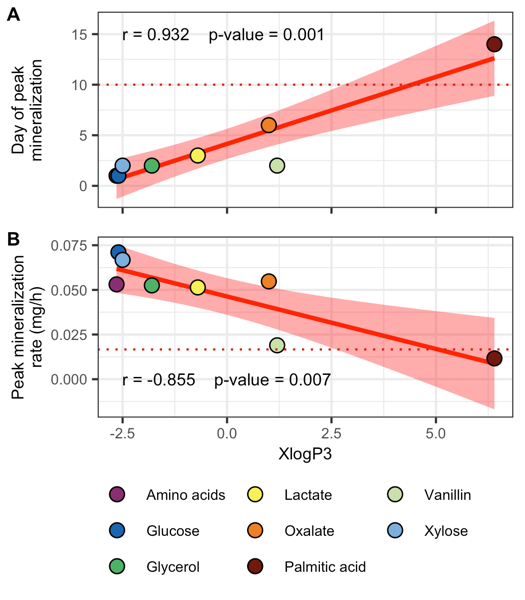

**Figure S3:** Predicted octanol-water partition coefficient (XlogP3) is significantly correlated to A) the day when maximum mineralization rate occurred and B) the maximum rate of substrate mineralization. Red lines represent linear regressions between factor and logP, with shading indicating the standard error. Pearson’s correlation analysis results are displayed in each plot.

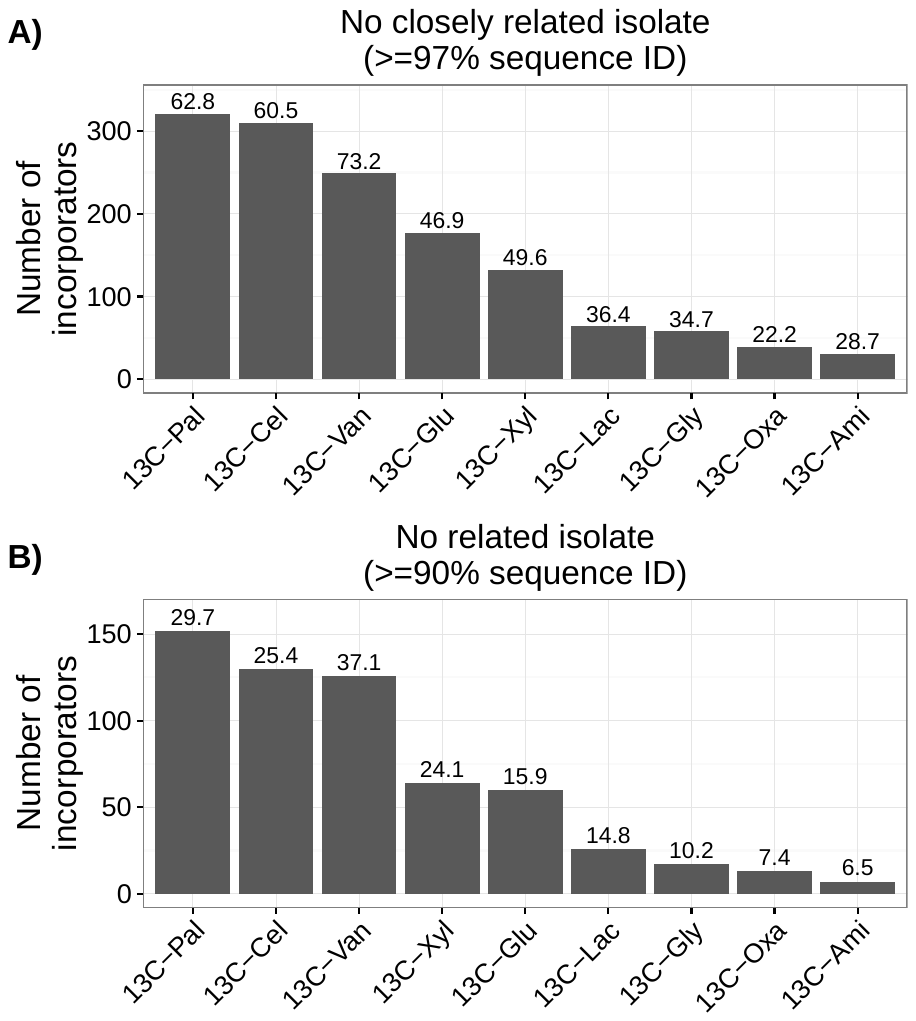

**Figure S4:** Many incorporators, particularly those assimilating C from C sources with low bioavailability, have no closely related isolates and in many cases, no related isolates. Represented here are the number of incorporators of each C source with A) no closely related isolates (≥ 97% sequence identity) and B) with no related isolates (≥ 90% sequence identity). Relatedness was determined from BLASTn queries of the OTU representative 16S rRNA gene sequences (V4 region) against sequences in “The All-Species Living Tree” project. Values above the bars are expressed as a percentage of the total number of incorporators for each C source.

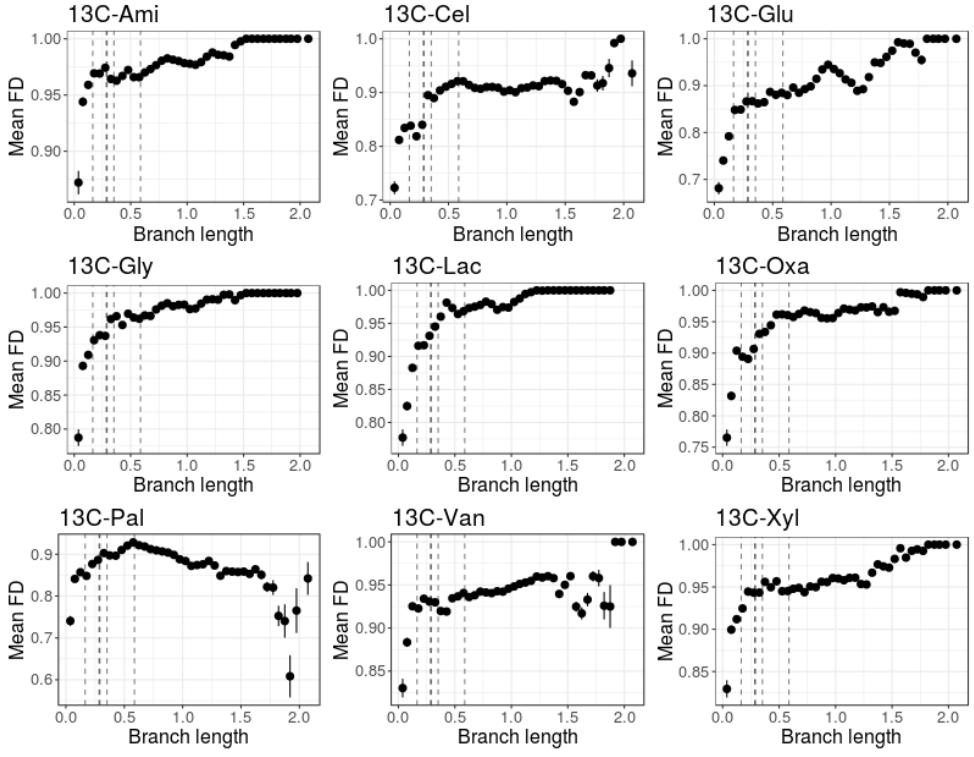

**Figure S5:** Phylogenetic distance (branch length) is a poor predictor of functional distance (mean FD) for incorporators when considering each C source independently. Branch length was derived from the 16S rRNA gene V4 region sequence phylogeny of all incorporators. Dashed lines are the median branch lengths separating OTUs across taxonomic groupings (left to right: genus, family, order, class, phylum).

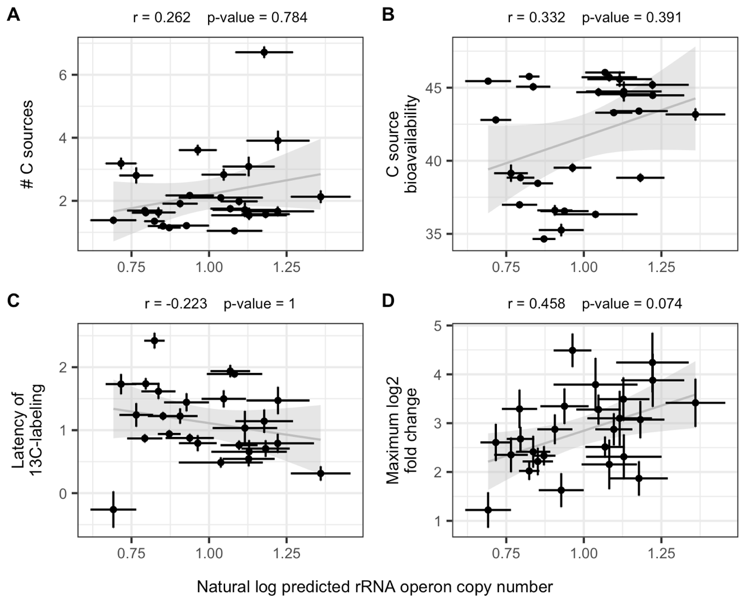

**Figure S6:** After removing the two AC with the highest predicted rRNA operon copy number (AC 1 and AC 19), the relationship between the natural log of the predicted rRNA operon copy number, a life history trait associated with the copiotroph-oligotroph continuum and four C assimilation characteristics: A) The average number of C sources from which C was assimilated by each guild. B) The averaged bioavailability of the sources from which C was assimilated by each guild. Bioavailability of a C source was defined as 48 (*i.e.,* experiment length) minus the day of peak C mineralization for that substrate. C) The averaged latency of C assimilation for each guild. D) The dynamic growth response of each guild as measured by increase in abundance in response to substrate addition. Red and grey lines indicate the statistically significant and non-significant linear relationship between factors respectively with shading representing standard error. Pearson’s r and p-values for these relationships are listed above each plot. p-values were corrected for multiple comparisons with Benjamini-Hochberg procedure (*n* = 4). Error bars for points indicate ± standard error across OTUs within each guild.

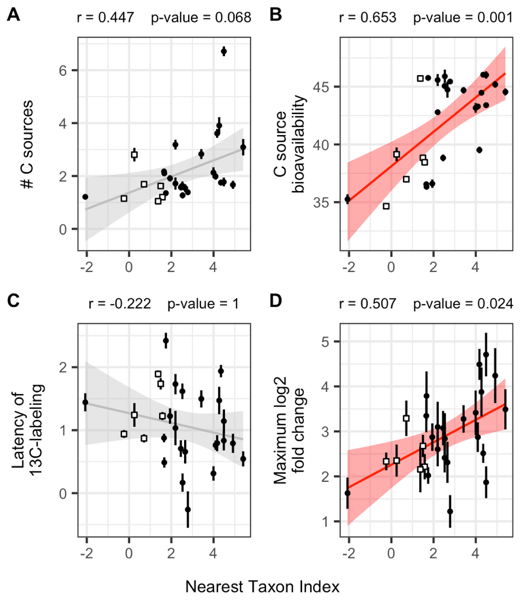

**Figure S7:** We tested whether the nearest taxon index within guilds was correlated with C assimilation and growth characteristics. A) The average number of C sources from which C was assimilated by each guild. B) The averaged bioavailability of the sources from which C was assimilated by each guild. Bioavailability of a C source was defined as 48 (*i.e.,* experiment length) minus the day of peak C mineralization for that substrate. C) The averaged latency of C assimilation for each guild. D) The dynamic growth response of each guild as measured by increase in abundance in response to substrate addition. Red and grey lines indicate the statistically significant and non-significant linear relationship between factors respectively with shading representing standard error. Pearson’s r and p-values for these relationships are listed above each plot. p-values were corrected for multiple comparisons with Benjamini-Hochberg procedure (*n* = 4). Error bars for points indicate ± standard error across OTUs within each guild.

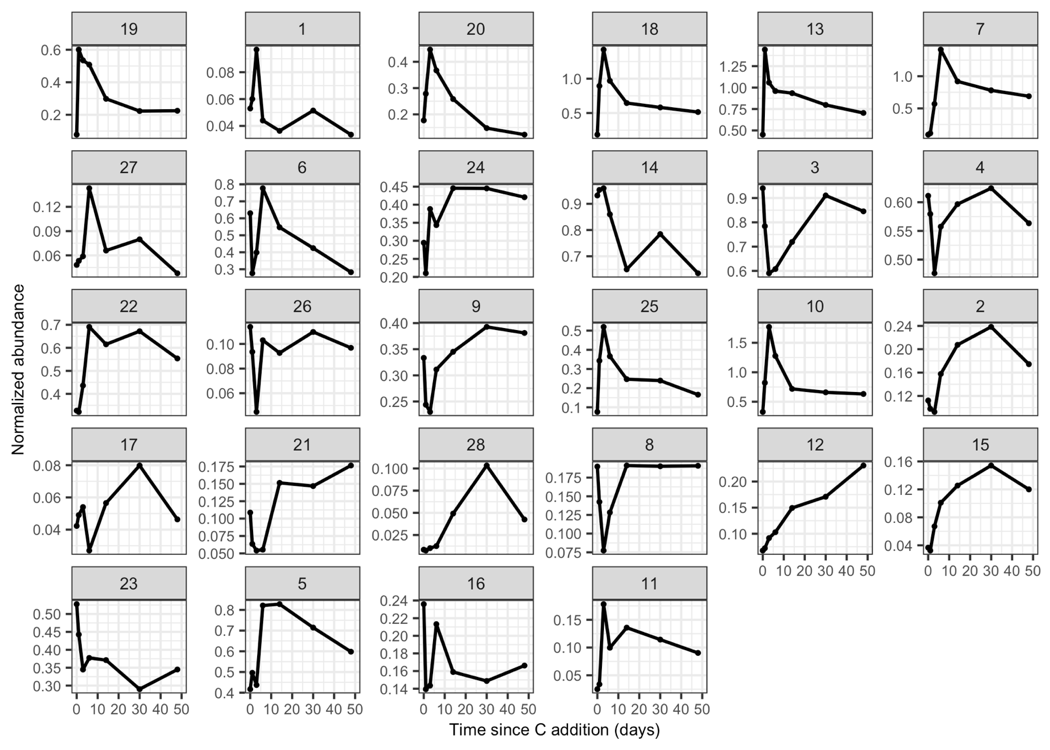

**Figure S8:** Normalized abundance of each guild over time throughout the experiment. Abundances are normalized by *rrn* copy number and DNA yield from the microcosms and then summed across all OTUs within each guild. Guilds are indicated by the number at the top of the plots and ordered based on positions in the PCA.

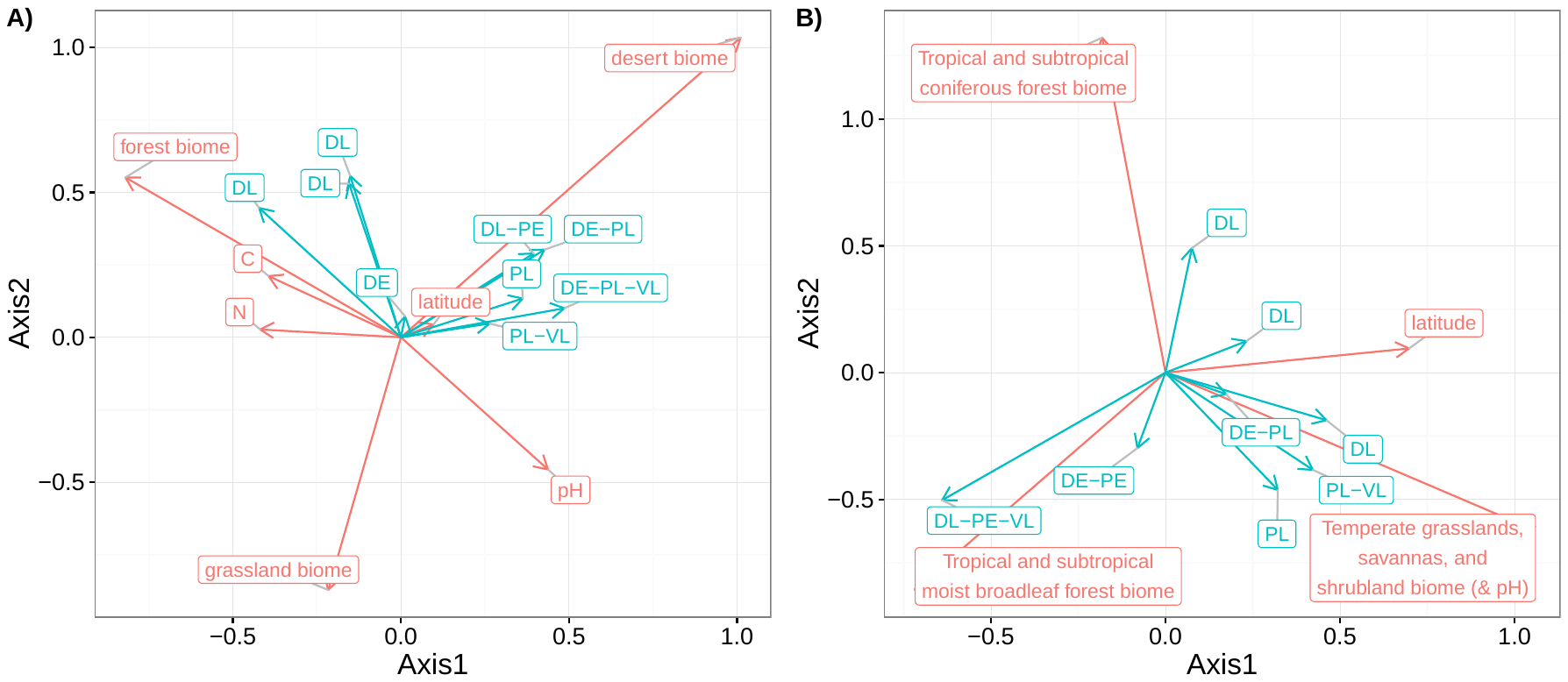

**Figure S9:** Results from RLQ analysis of incorporators mapped to bacterial surveys at the A) continental scale (QIITA study 619) and B) global scale (QIITA study 928). Incorporator functional clusters were further grouped by form of the substrate from which C was assimilated (D = dissolved, including glucose, xylose, amino acids, glycerol, lactate, and oxalate; V = vanillin; P = particulate, including cellulose and palmitic acid) and by time of C assimilation (E = early, L = late) as indicated by the blue letters.
